# Integer topological defects provide a new way to quantify and classify cell sheets

**DOI:** 10.1101/2024.08.28.610106

**Authors:** Zihui Zhao, He Li, Yisong Yao, Yongfeng Zhao, Francesca Serra, Kyogo Kawaguchi, Hepeng Zhang, Hugues Chaté, Masaki Sano

**Affiliations:** School of Physics and Astronomy, Shanghai Jiao Tong University, Shanghai, 200240, China; Institute of Natural Sciences, Shanghai Jiao Tong University, Shanghai, 200240, China; Center for Soft Condensed Matter Physics and Interdisciplinary Research, Soochow University, Suzhou, 215006, China; Physics and Astronomy, Johns Hopkins University, Baltimore, 21218, USA; Department of Physics, Chemistry and Pharmacy, University of Southern Denmark, Odense, 113-0033, Denmark; Universal Biology Institute, The University of Tokyo, Tokyo, 113-0033, Japan; Nonequilibrium Physics of Living Matter RIKEN Hakubi Research Team, RIKEN Center for Biosystems Dynamics Research, Kobe, 650-0047, Japan; RIKEN Cluster for Pioneering Research, Kobe, 650-0047, Japan; Institute for Physics of Intelligence, The University of Tokyo, Tokyo, 113-0033, Japan; Service de Physique de l’Etat Condensé, CEA, CNRS Université Paris-Saclay, CEA-Saclay, Gif-sur-Yvette, 91191, France; Computational Science Research Center, Beijing, 100094, China

## Abstract

Sheets of confluent cells are often considered as active nematics, with accumulation at 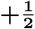 topological defects and escape from 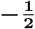 defects being widely recognized. However, collective dynamics surrounding integer-charge defects remain poorly understood, despite its biological importance. By using microfabricated patterns, we induce diverse **+1** topological defects (aster, spirals, and target) within monolayers of neural progenitor cells. Remarkably, cells are consistently attracted to the core of **+1** defects regardless of their type, challenging existing theories and the conventional extensile/contractile dichotomy. We trace back the origin of this accumulation behavior to previously overlooked nonlinear active forces using a combination of experiments and a continuous theory derived from a cell-level model. Our findings demonstrate that **+1** topological defects can reveal key features of active nematic systems and offer a new way to characterize and classify cell layers.

## 1 Introduction

Elongated cells forming confluent layers have long been described as liquid crystal nematics, passive [1] or, more recently, active [2–6]. These active nematics display half-integer topological defects, as in the passive case, where they are energetically more favorable than integer defects [7–15]. Recently gathered evidence shows that 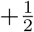 defects can be the location of cell accumulation or cell extrusion, whereas cell density is depleted near 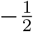 defects [16, 17]. This was in particular observed in confluent layers of Neural Progenitor Cells (NPCs). A simple hydrodynamic linear theory resting on the classic description of extensile active nematics was shown to account qualitatively for these findings [12, 16].

Integer topological defects are also prevalent in nature and biologically relevant, such as asters in a neural rosette, targets in plants meristem, and asters and targets during the formation of mouth, tentacles, and foot of hydra [18–22]. Integer defects do not appear spontaneously in cellular nematic layers, as they tend to split into pairs of half-integer ones. However, such defects could be induced by external factors and thus also play crucial biological functions. Recently, in vitro studies have shown that +1 defects induced by strong circular confinement provide sites for growth and cellular differentiation [23]. These C2C12 myoblast cells exhibit nematic order in wider space [16]. Still, confinement leads to polar order, accumulation at defect centers and inward flow, but with a small system size and non-negligible boundary effects. Despite the biological importance of morphogenesis, we still lack quantitative measurements of tissue properties and large-scale flow dynamics of active nematics within stable and well-defined topological defect configurations.

In dense conditions, NPCs take elongated shapes and display stochastic back-and-forth motion without apparent contact inhibition or strong cell-cell adhesion [16], exhibiting local nematic order. These characteristics make NPCs an excellent system for building hierarchical descriptions of active nematics, from the individual cell level to continuous theory, a rare feat we take advantage of here.

To investigate the behavior of cellular active nematics with integer defects, we induce +1 defects in NPC layers by letting cells proliferate on patterned substrates. We consider a one-parameter family of +1 defects from aster to target via spiral vortices and show that cells accumulate at their center in all cases, suggesting potential biological importance. A similar effect was recently observed with fibroblasts proliferating on a target pattern [24, 25], and attributed to a higher cell division rate at the defect core rather than to the large-scale motion of cells typical of hydrodynamic theories [25]. In contrast, we demonstrate here that our system does exhibit large-scale inward cellular flows. We further show that our findings cannot be explained by the simple hydrodynamic theory used in the unpatterned ‘natural’ case, since this theory predicts outward flows for the target. We perform a complete modeling of our system, starting from a minimal particle/cell level model incorporating its basic features, deriving a nonlinear hydrodynamic theory from it, and analyzing its solutions. Details of this modeling program will appear elsewhere [26]. Analysis of the experiment, based on this theory, reveals that nonlinear active force terms neglected in simpler approaches play a crucial part in explaining cellular flows around defects. We estimate all the coefficients that characterize the theory. Our results suggest a new classification of active nematics and reveal important physical mechanisms heretofore ignored, which are likely to play a role in various contexts given the genericity of our approach.

## 2 Results

### 2.1 NPCs accumulate at the center of +1 topological defects

To investigate the effect of +1 topological defects on cell monolayers, we cultured NPCs on microfabricated poly-dimethylsiloxane (PDMS) patterns featuring various defect types delineated by 1.2*μm*-high ridges (Fig. 1a, Methods). These patterns are defined by their tilt angle *θ*_0_: everywhere in space, but here only along the fabricated shallow ridges, *θ*_*p*_ = *ϕ* + *θ*_0_, where *θ*_*p*_ is the preset local orientation, *ϕ* is the polar angle, and *θ*_0_ (*θ*_0_ ∈ (−*π/*2, *π/*2]) is the tilt angle relative to the radial direction (Fig. 1b) [27]. Specific values *θ*_0_ = 0, ±*π/*12, ±*π/*6, ±*π/*4, ±*π/*3, ±5*π/*12, *π/*2 were used. Note that *θ*_0_ = 0, 0 *< θ*_0_ *< π/*2, −*π/*2 *< θ*_0_ *<* 0, *θ*_0_ = *π/*2, correspond to aster, counterclockwise spirals, clockwise spirals and target respectively (Fig. 1c). The patterns had a maximal radius *R*_*max*_ = 500*μm*, and included a central ridge-less area with a radius *R*_*min*_ = 80*μm*. The shallow ridges can be considered to constitute weak external fields since the cells retained their ability to easily climb and cross over them (Fig. 1d, Supplementary Video 1). They gently guided cells and led to the emergence of +1 defects in the confluent cell monolayer (Fig. 1f).

**Fig. 1.**
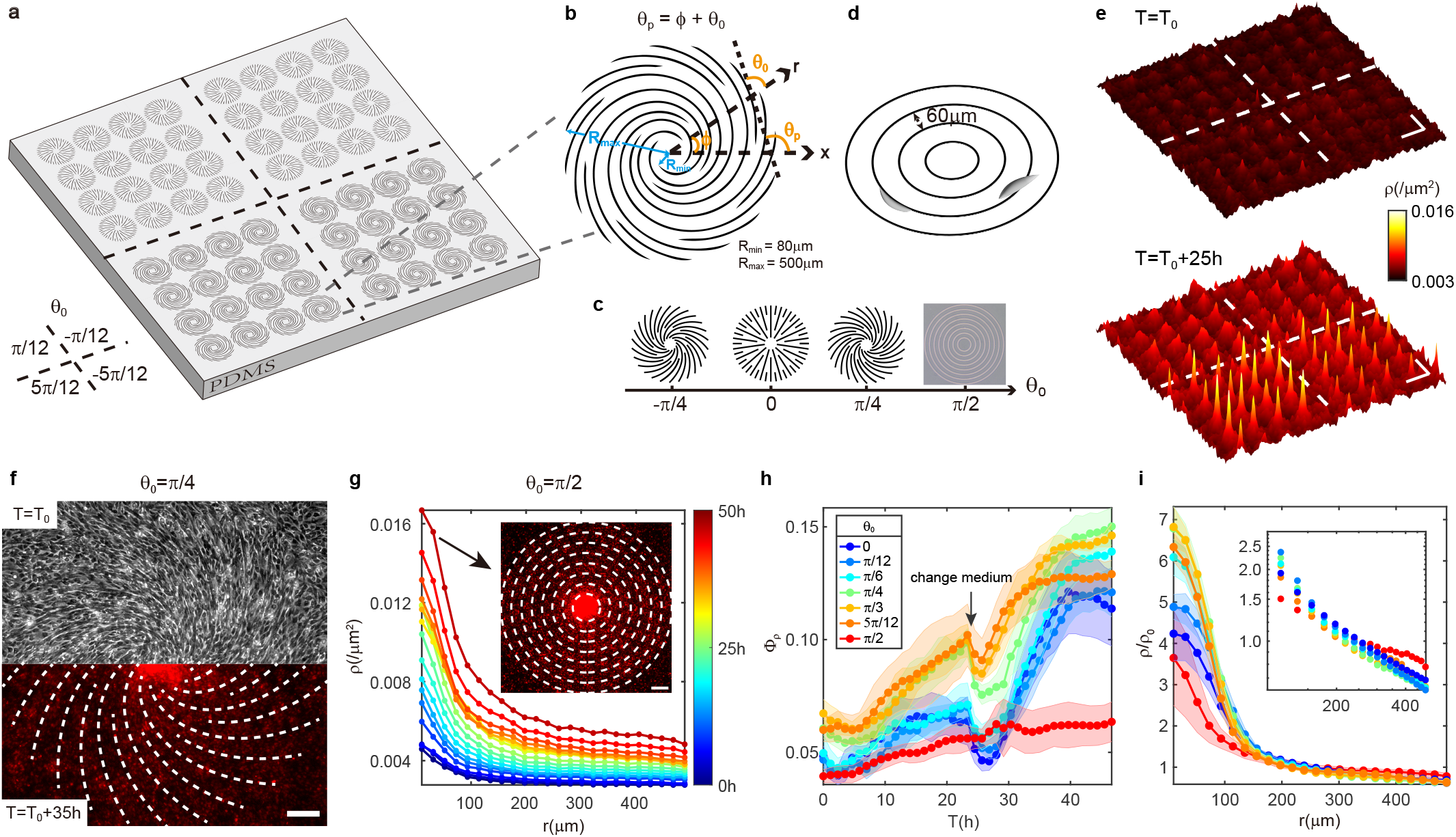
NPCs accumulate at centers of asters, spirals, and targets. **a**, Schematic of microfabricated patterns on a PDMS substrate. Four arrays of different spirals (*θ*_0_ = ±*π/*12, ±5*π/*12) are arranged on one PDMS sheet, featuring 16 micropatterns per defect type. **b**, Schematic of +1 topological defect definition. The largest radius of patterns (*R*_*max*_) is 500*μm*. The radius of the defect center without ridges (*R*_*min*_) is 80*μm*. The distance between ridges is in the range of 30 − 120*μm*. **c**, Types of +1 topological defects characterized by the tilt angle *θ*_0_. A bright-field image is shown for targets. **d**, Schematic of cell motion on a target pattern. **e**, Heatmap of cell density at the beginning and after 25 hours of recording on spirals shown in **a**. *T* = *T*_0_ at the onset of confluence. **f**, A typical phase-contrast image at the beginning and a typical fluorescent image after 35 hours of recording on a spiral with *θ*_0_ = *π/*4. Dashed lines depict ridges of patterns. **g**, Time evolution of average radial density profiles on the target. The data was averaged every one hour and by 32 targets. Inset: a typical fluorescent image showing accumulation after 50 hours of recording on a target. Dashed lines depict ridges. **h**, Average percentage of cells Φ_*ρ*_ (Φ_*ρ*_ = *n*(*r < R*_*min*_)*/n*(*r < R*_*max*_), where n is the cell number) assemblies over time on aster, counterclockwise spirals and target. Colors depict different defects. The number (N) of each type of defect is 6 and all defects was designed on one PDMS sheet. **i**, Normalized radial density profiles by mean density *ρ*_0_ when *ρ*_0_ = 0.009*/μm*^2^. Inset: A log-log plot for the region with *r >* 100*μm*. N of aster, each type of spiral, target is 32, 6 and 32. Legend in **h**. In **h** and **i**, data are presented as mean ± s.d., and s.d. are calculated from defects in an array. Scale bars, 100*μm*.

We conducted long time-lapse recordings after confluence and observed that cells gradually accumulated over time at the centers of defects irrespective of *θ*_0_ (Fig. 1e,f,g, Methods). The local cell density *ρ* was measured from nuclear fluorescence intensity (Methods). Averaging the configurations azimuthally, we obtained the time-evolution of cell density radial profiles across all patterns (Fig. 1g, Extended Data Fig. 1a,2a). The ratio Φ_*ρ*_, representing the number of cells within the radius *R*_*min*_ around defect center relative to the total number, exhibits a continuous increase except during periods of medium changes (Fig. 1h, Extended Data Fig. 1b,2b, Supplementary Video 2) [28]. Radius-dependent average cell densities on different defects with the same total density display distinct accumulation around centers and similar slopes in log-log plot in the regions other than centers (Fig. 1i, Extended Data Fig. 1c). Additionally, the accumulation behavior is more prominent on counterclockwise patterns, potentially attributable to NPCs’ weak chirality (Extended Data Fig. 2b,2c) [29].

### 2.2 Orientation and dynamics of NPCs in induced topological defects

When forming a confluent monolayer, cells demonstrated a tendency to align with their neighbors and with the guiding patterns (Fig. 2a-c). The local orientation of this nematic order hardly changed throughout the recording. Applying the structure tensor method to phase-contrast images and averaging orientations locally over space and time, we obtained local nematic orientation fields (Fig. 2a-c, Extended Data Fig. 3a, Methods). To quantify the degree of orientational order, we calculated the local scalar order parameter *S. S* is minimal at the center, increases with radius, and slightly decreases near the edges due to the looser confinements (Fig. 2d, Extended Data Fig. 3d). However accurate orientation measurements were hindered around defect centers due to the presence of high densities and three-dimensional cell mounds in these areas. Coherency analysis also indicates that the estimated orientations are more reliable in outer regions than around centers. (Extended Data Fig. 3e,f, Methods).

**Fig. 2.**
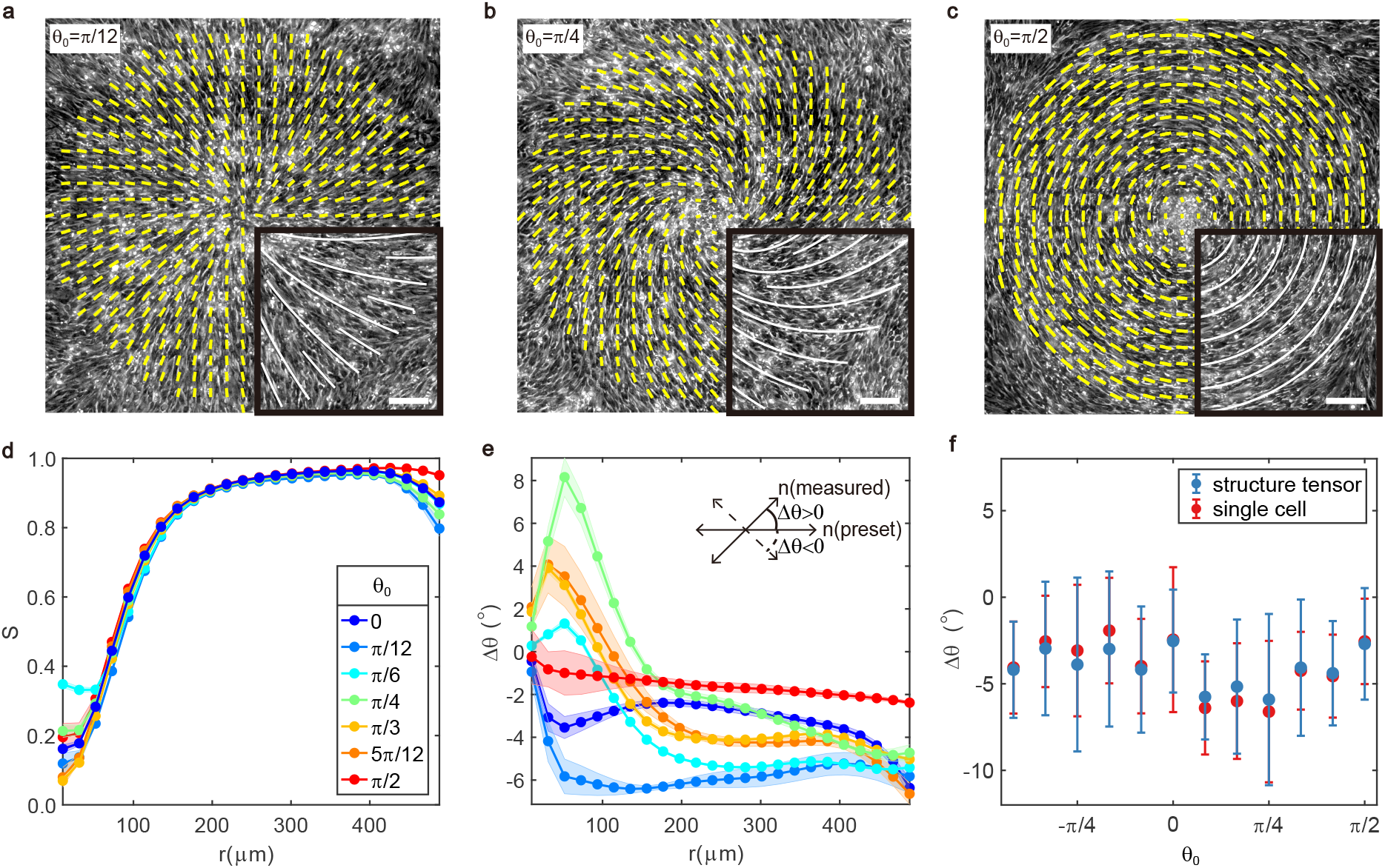
Cellular orientation around induced +1 defects. **a, b, c**, Spatiotemporally averaged orientational fields on defect pattern with *θ*_0_ = *π/*12, *π/*4, *π/*2, respectively. Yellow lines depict local nematic orientation, with their length corresponding to local coherency. Background: phase-contrast images, from which we can observe cell bodies. Insets: white lines show the shallow ridges defining patterns. **d**, Radial profiles of order parameter *S* on aster, counterclockwise spirals and target. **e**, Same as **d** but for the angle Δ*θ* between the experimentally measured cell orientation and orientation preset by PDMS pattern. Preset orientation refer to cellular alignment if cells form perfect topological defects according to definition. Inset shows the definition of Δ*θ*: Δ*θ >* 0 when the measured orientation (measured **n** = (cos *θ*, sin *θ*)) rotates counterclockwise against the preset orientation (preset **n** = (cos *θ*_*p*_, sin *θ*_*p*_)), vice versa Δ*θ <* 0. **f**, Δ*θ* across all types of defects. In **a**−**f**, data are averaged by 32 asters (10h), 16 spirals for each type (10h) and 32 targets (10h). In **d, e**, data are presented as mean ± s.d., and s.d. are calculated from different times. In **f**, data are presented as mean ± s.d., and s.d. are calculated from different positions of average orientational fields (regions with *r <* 100*μm* are excluded). Scale bars, 100*μm*.

To evaluate the impact of guiding ridges, we measured the angle Δ*θ* between local cell orientation (*θ*) and the preset pattern orientation (*θ*_*p*_). We found that Δ*θ* does not change much over radius (Fig. 2e, Extended Data Fig. 3g). To avoid inaccuracies, we did global average in regions with *r >* 100*μm* and consistently observed that Δ*θ* was around −4^°^ across all defects, indicating a weak clockwise chirality (Fig. 2f). Comparable results were obtained from orientational fields extracted from the long axis of nuclei (Fig. 2f, Extended Data Fig. 3b,c, Methods). This observation of the small value validates the successful induction of the designed defects in the cell monolayer.

To investigate whether the aggregation in defect central areas is related to collective movements, we conducted a quantitative analysis of cell dynamics. Trajectories depicted in (Fig. 3a-b) were extracted by tracking cell nuclei (Methods, Supplementary Video 3-6). Individual cells exhibit bidirectional motion with an instantaneous speed around 0.4*μm/min* and a typical reversal time of 1-2 hours (Fig. 3c, Methods). By coarse-graining instantaneous velocities during the continuous accumulation periods (10 hours) in Fig. 1h, we obtained spatiotemporally averaged net velocity fields (Extended Data Fig. 4a, Methods, see Supplementary Video 3-6 for time evolution of spatially averaged net velocity fields). Streamlines calculated from these net velocity fields demonstrate the inward direction of cellular flows (Fig. 3d-f). Notably, the local velocities are typically not aligned with the local orientation. Averaging radial net velocities along the azimuthal direction shows inward net radial velocities across all defects (Fig. 3g, Extended Data Fig. 4b), consistent with observed cell accumulation around defect cores. In our measurements here and subsequent fitting process, we focused on a timescale of 10 hours, during which cells did not proliferate much since cell division typically occurs over approximately 24 hours [16].

**Fig. 3.**
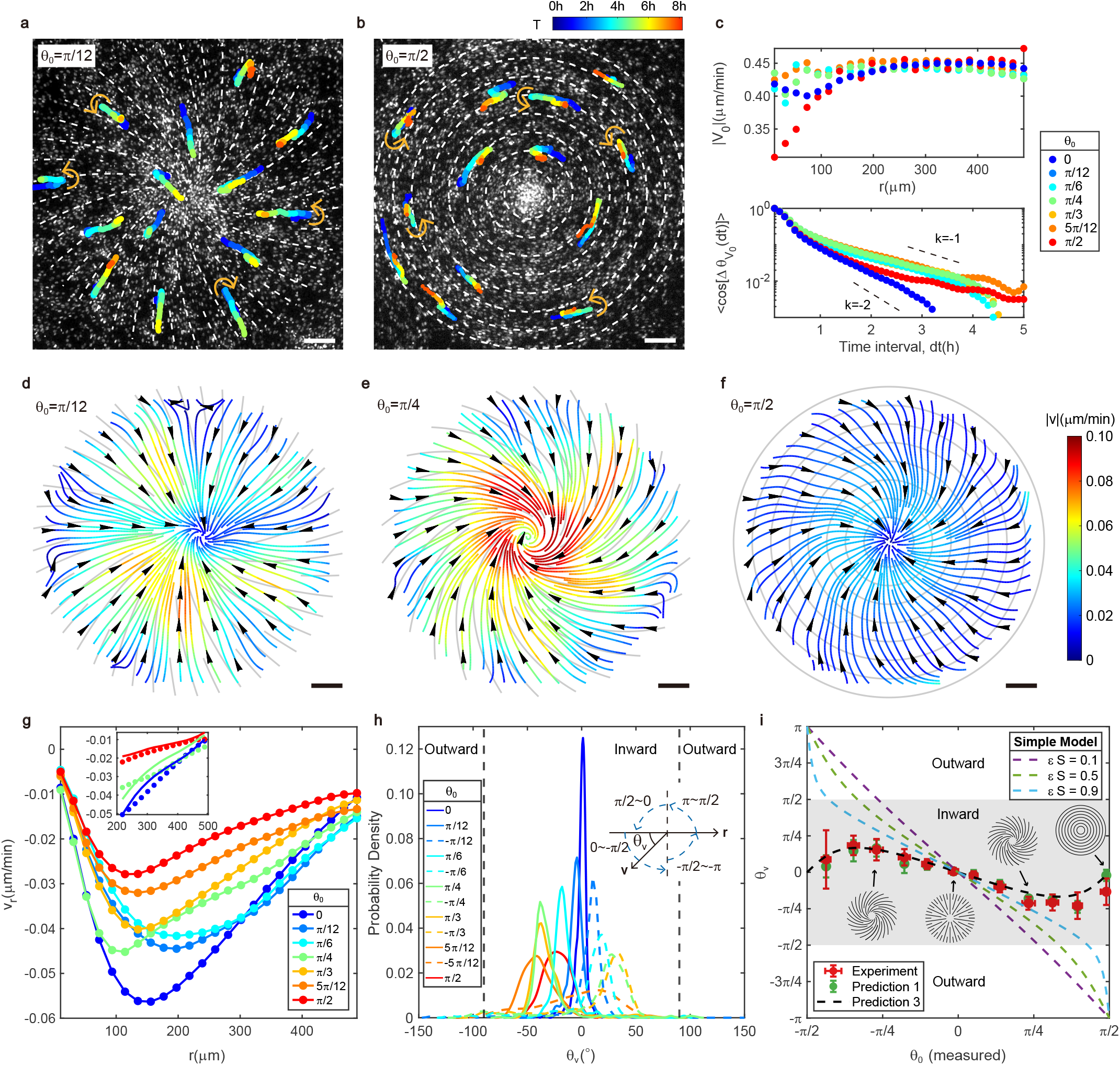
Cellular dynamics during accumulation toward +1 defect cores. **a, b**, Bidirectional motion of single NPC on defects with *θ*_0_ = *π/*12, *π/*2, respectively. Trajectories are traveled in time as indicated by color. Color bar in **b**. Background: fluorescent images, from which we can observe cell nuclei. Dashed lines indicate the guiding ridges of patterns. **c**, Radial profiles of the amplitude of average instantaneous velocity *V*_0_ when mean density is around 0.005*/μm*^2^ (upper) and autocorrelation of the direction 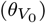 of instantaneous velocity as a function of time interval *dt* (lower). *k* = −1 (*k* = −2) corresponds to 2 hours’ (1 hour’s) reversal time. **d, e, f**, Streamlines on defects with *θ*_0_ = *π/*12, *π/*4, *π/*2, respectively. Gray lines indicate guiding ridges. Color represents the amplitude of net velocities. Color bar in **f**. **g**, Radial profiles of the average radial net velocity (*V*_*r*_) components on aster, counterclockwise spirals and target. Inset: Comparison between experimentally measured radial velocity profile (dots) and predictions (lines) at large r on the aster, spiral (*θ*_0_ = *π/*4) and target. The prediction comes from method (ii). **h**, Probability density function of the angle (*θ*_*v*_) between net velocity and inward radial direction on all types of defects. Insets show the definition of *θ*_*v*_. **i**, Measured and predicted *θ*_*v*_ versus measured *θ*_0_. *θ*_0_(*measured*) = *θ*_0_ + Δ*θ*. Prediction 1 (3) comes from method (i) ((iii)). Data are presented as mean ± s.d. in horizontal axis and the location of peak ± FWHM of PDF shown in **h** in vertical axis. FWHM and s.d. are calculated from different positions of average net velocity fields (regions with *r <* 100*μm* are excluded). In **c** − **i**, data are averaged by 32 asters (10h), 16 spirals for each type (10h) and 32 targets (10h). Scale bars, 100*μm*.

We further analyzed the probability distribution function of angles (*θ*_*v*_) between net velocities and inward radial directions. To avoid inaccuracies of tracking single cells in high-density centers, we excluded the regions with *r <* 100*μm*. These angles (*θ*_*v*_) are predominantly situated in the inward region (−90^°^ *< θ*_*v*_ *<* 90^°^) (Fig. 3h). After extracting the peak location of *θ*_*v*_ for each defect in (Fig. 3h) and incorporating the deviations Δ*θ* in (Fig. 2f) into preset *θ*_0_ (*θ*_0_(*measured*) = *θ*_0_ + Δ*θ*), we finally delineated how *θ*_*v*_ changes with measured *θ*_0_ (Fig. 3i). Comparable results were obtained with optical flow method (Extended Data Fig. 4c,d, Methods). Symmetry-breaking shown in Fig. 2f, Fig. 3f, Fig. 3h indicates the presence of chirality.

### 2.3 Detecting the relation between net cellular flows and active forces

Conventional active nematic theories typically include an equation relating a velocity field to an active force term:

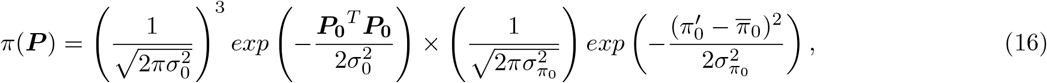

where *γ* is friction coefficient, and 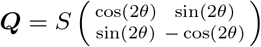 is the nematic order parameter tensor with *θ* being the local cell orientation. The active force (***f***) can formally be identified as the divergence of active stress −*ζ****Q***, leading to ***f*** = −*ζ*∇ · ***Q*** [30, 31]. In the ‘unpatterned’ natural state, 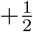 defects in the confluent NPCs are observed to move in the ‘head’ direction [16, 32], considered as a characteristic of an extensile system withζ *>* 0.

In our experiments, cells aligned well with the guiding patterns, resulting in a significantly slower timescale for ***Q*** dynamics compared to accumulation processes. Therefore, we assume ***Q*** is fixed in time by the ‘external field’ induced by the pattern. For +1 defects with *θ* = *θ*_*p*_ = *ϕ* + *θ*_0_, the linear active force field becomes 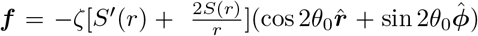, where 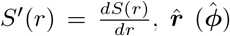 is the unit vector in the radial (azimuthal) direction. In a uniform and confluent state where *S*^*′*^(*r*) *>* 0 near the center, the radial component is inward for 0 ≤ |*θ*_0_| *< π/*4 but outward for *π/*4 *<* |*θ*_0_| ≤ *π/*2, indicating possible outward flows for large |*θ*_0_| (Fig. 4a). This is at odds with the inward flows observed with all types of induced +1 defects. In particular, this suggests that within this simple theory, the collective behavior around large |*θ*_0_| defects like targets is that of a contractile system (*ζ <* 0).

**Fig. 4.**
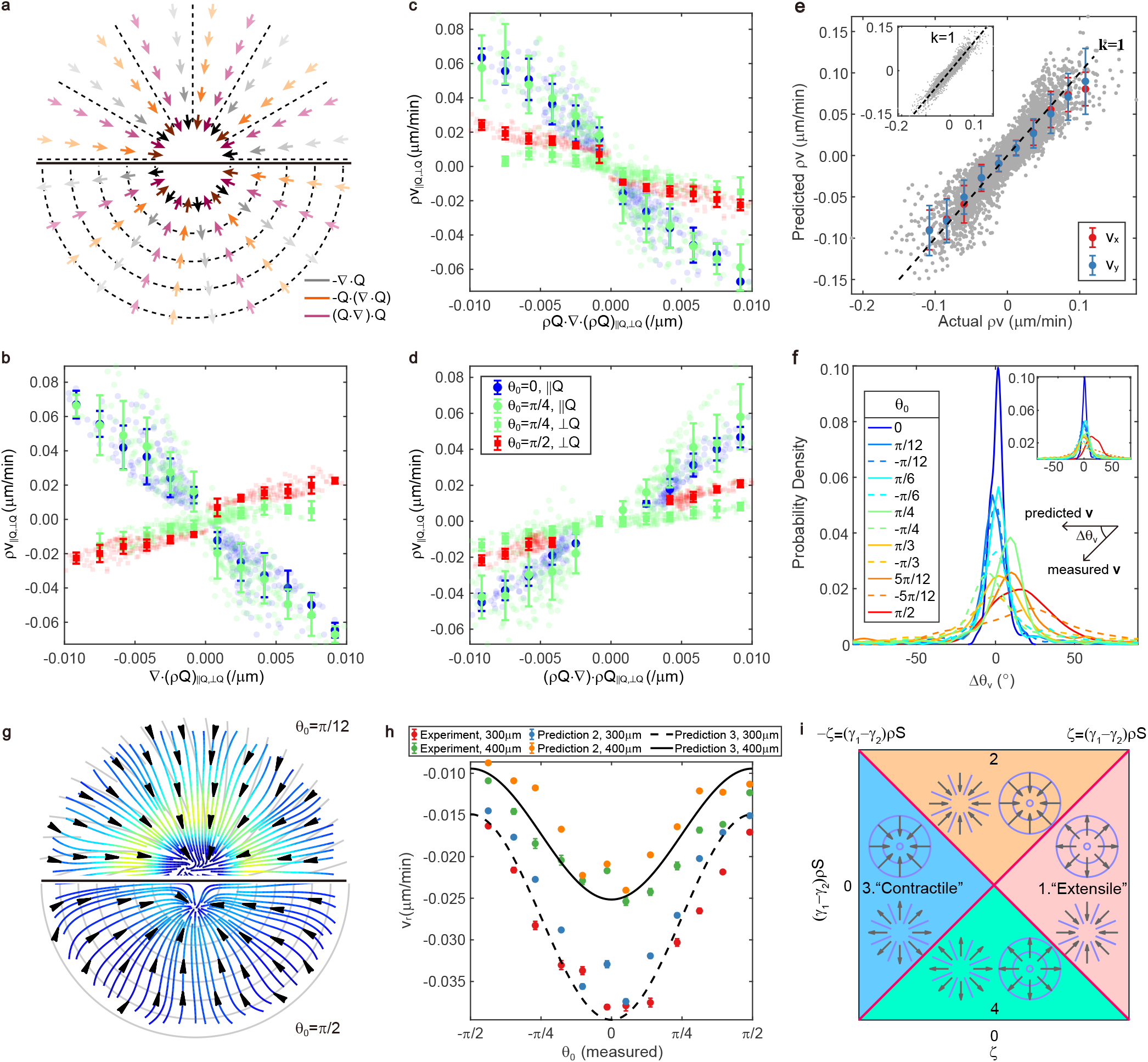
Prediction of cellular flows from active forces. **a**, Schematic of active forces on aster and target when *S* is a constant. Darker color means stronger force. **b, c, d**, Comparisons between components of cellular flow *ρv* and components of active forces ∇ · (*ρ****Q***) (**b**), *ρ****Q***·(∇·*ρ****Q***) (**c**), (*ρ****Q***·∇)·*ρ****Q*** (**d**). *ρv*_∥***Q***_ (*ρv*_⊥***Q***_) is defined as the flow component parallel (perpendicular) to the principal axis of ***Q***, so as for components of active forces. Comparisons between parallel components are plotted for the aster and spiral (*θ*_0_ = *π/*4); Comparisons between perpendicular components are plotted for the target and spiral (*θ*_0_ = *π/*4). Error bars are s.d. **e**, Comparison between experimentally measured cellular flows and predictions. Determinate coefficient *R*^2^ = 0.8173. The scatter plot denotes raw data (gray) on all the patterns (regions with *r <* 100*μm* are excluded). Symbols show the average values of predictions. The prediction method is method (ii). Error bars are s.d. Insets show results from prediction method (i). In **b** − **e**, *ρ* is normalized by mean density. **f**, Probability density function of the angle (Δ*θ*_*v*_) between experimentally measured net velocities and predicted net velocities. The prediction method is method (ii). Insets show the definition of Δ*θ*_*v*_ and results from prediction method (i). **g**, Predicted streamlines on the spiral (*θ*_0_ = *π/*12) and target. Color bar in Fig. 3**f**. The prediction method is method (i). **h**, Comparison between measured net radial velocity and predictions at *r* = 300*μm* and *r* = 400*μm* of all the defects. Method of prediction 2 (3) is method (ii) ((iii)). Data are presented as mean ± s.e.m. **i**, New classification of active nematics. The functions defining two red lines are shown on the top left and top right.

The linear theory can be refined by considering an anisotropic friction *γ* = *γ*_0_(*I* −*ϵ****Q***), with the anisotropy coefficient 0 ≤ *ϵ* ≤ 1 describing larger friction along the direction perpendicular to cell alignment [12, 16]. However, comparing measured *θ*_*v*_ with predicted *θ*_*v*_ from this extended simple theory shows that this fails to explain cell accumulation and inward flows for all defects types (Fig. 3i).

To understand the underlying mechanisms driving inward cellular flows, we performed a complete modeling of our system, starting from a minimal particle/cell level dry active nematic model where the induced patterns are accounted for by some external field. Numerical simulations of this model showed particle accumulation irrespective of the type of +1 defect, in broad agreement with our experimental results. Confident in the faithfulness of this model, we derived from it a nonlinear hydrodynamic theory using a Boltzmann-Ginzburg-Landau approach (Supplementary Information: Section 4) [26, 33–35]. The theory shows the presence of two ‘nonlinear active force terms’ in addition to the standard active force. Incorporating these terms and accounting for spatial density variations, the total active force replacing ***f*** is

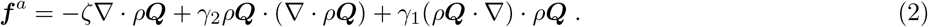

In the steady state with ideal orientations, *θ* = *θ*_*p*_ = *ϕ* + *θ*_0_, equation (2) can be written in polar coordinates as

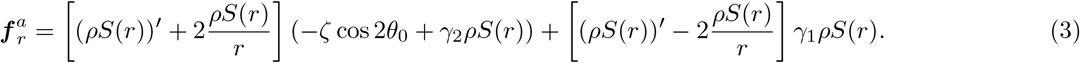

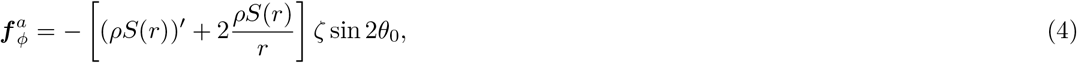

where 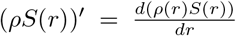. Our derivation shows that *γ*_2_ *<* 0 and that *γ*_1_ *>* 0 generally holds in most cases (Supplementary Information: Section 4). Therefore, the two nonlinear active forces point towards the defect cores, which contributes to inward flows and central cell accumulation (Fig. 4a) (Note that a ***Q*** · (∇· ***Q***) term was considered in [36] in the context of active suspensions, where it was shown to be able to stabilize nematic order). Our theory taking into account variations of the cell density *ρ*, the net velocity of cellular flow is governed by

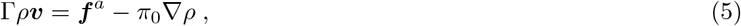

where Γ= *γ*_0_(*I* − *ϵρ****Q***), 0 ≤ *ϵρ* ≤ 1 and *π*_0_ *>* 0.

To test the relation between cellular flow and active forces from experimental data, we decomposed them into the parallel and perpendicular components relative to the principal axis of local ***Q*** in regions outside centers (Extended Data Fig. 5). In Fig. 4b-d, we present parallel components on asters and spirals (*θ*_0_ = *π/*4) and perpendicular components on targets and spirals (*θ*_0_ = *π/*4) as they represent the primary components associated with inward flows on corresponding patterns. Concerning the ∇· *ρ****Q*** term, the parallel components exhibit negative correlations with the net velocity, while perpendicular components show positive ones (Fig. 4b), another indication that the simple theory cannot fully explain the observed phenomena. However, for *ρ****Q***·(∇·*ρ****Q***) and (*ρ****Q***·∇)·*ρ****Q*** terms, these components show consistent negative (Fig. 4c) and consistent positive correlations (Fig. 4d) respectively. These comparisons provide evidence for the relevance of the two nonlinear active forces.

### 2.4 sPredicting cellular flows from local orientations

From experimental data, we extracted the value of each term in equations (2,5) using the average net velocities, orientations, and densities outside the centers. To quantitatively assess our theory, we of course also need to estimate their coefficients. We identified the best-fit values by Bayesian inference. During fitting, we replaced *ρ* in equations (2,5) with the normalized density *ρ/ρ*_0_, where *ρ*_0_ is the mean density. The fits yielded values for five model parameters: 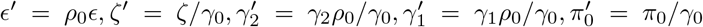. For each defect type, we determined the best-fit values (Extended Data Tab. 1). We then averaged these best-fit values over the different defect types and found: *ϵ*^*′*^ = 0.1476±0.0876,*ζ*^*′*^ = 0.0289±0.0131(*μm*^2^*/s*), 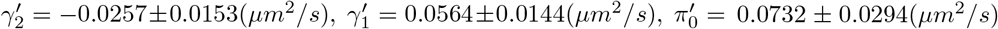, with standard deviations as errors. It is worth noting that since not all experiments were conducted simultaneously, standard deviations may be large due to varying cell conditions. However, the signs 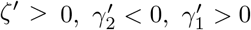 are maintained and the order of magnitude of the coefficients is found steady.

We conducted three distinct approaches to predict cellular flows from equation (5). (i) Utilize the measured ***Q*** and *ρ* fields from experiments for each defect type with best-fit parameters; (ii) Employ the same fields but use average parameters; (iii) Generate an *S*(*r*) profile and a *ρ*(*r*) profile by averaging across all defects, use average parameters and active force components in equations (3,4). By comparing the amplitudes and angles of measured and predicted net velocities, we discovered that while the best-fit parameters yielded the most accurate predictions, the average parameters also provided acceptable predictions (Fig. 4e,f). Through methods (i) and (ii), we obtained predicted velocity fields for all defects (Extended Data Fig. 6a,7a) and displayed streamlines on a spiral and target (Fig. 4g). In Extended Data Fig. 8 and Fig. 3g, we compared the predicted and measured net radial and azimuthal velocity profiles. Further, outside the centers, where density profiles of all defect types are similar, the measured and predicted net radial velocities exhibit the same cosine-like tendency over *θ*_0_ (Fig. 4h), consistent with the radial components of active forces in the equation (3). Moreover, the predicted tendency of *θ*_*v*_ over *θ*_0_ closely matched the measured tendency, contrasting sharply with the simple theory (Fig. 3i, Extended Data Fig. 6b,7b). In the prediction, we obtained slight outward flows at centers of defects with large |*θ*_0_| (Fig. 4g, Extended Data Fig. 8a,b). By using ideal orientations and setting 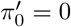 at central regions according to an assumption, the outward cellular vanished basically (Extended Data Fig. 9, Methods).

We finally propose a classification of dry active nematics based on our observed phenomena, given that our system cannot be categorized simply as contractile or extensile (as determined by the sign ofζ) due to the nonlinear active forces. Neglecting the gradients of *ρ* and *S* in equation (3), the direction of force along the radial direction is governed by the larger term between ±*ζ* and (*γ*_1_ − *γ*_2_)*ρS*, where + (−) is for target (aster). When the nonlinear active forces dominate the linear terms ((*γ*_1_ − *γ*_2_)*ρS >* ±*ζ*), two new cases (region 2 and 4 in Fig. 4i) appear, in which both target and aster can be attractive or depleted. In our case, NPC monolayers are situated in the region 2, which has never been classified before.

## 3 Discussion

In this work, we explored the dynamics of NPC monolayers with +1 topological defects. Using shallow-ridged substrates to induce aster, spirals, and target, we observed cell accumulation near their center and inward cellular flow towards defect cores, regardless of the specific defect type. Analysis of experimental data, including orientations, velocities, and densities, revealed the origin of cellular flows across various defects, stressing the existence of the nonlinear active forces suggested by the hydrodynamic theory we derived from a relevant particle-level model. By incorporating these active forces, we quantitatively reproduced cellular flows in large-scale regions.

We believe that the accumulation phenomena described here are essentially due to generic properties of dense 2D cellular active nematics, despite some measurements of cell mounds at centers of defects being inaccurate. The chiral symmetry-breaking witnessed at various points along this work (Fig. 2f, Fig. 3f,h, Fig. 4h, Extended Data Fig. 2c) warrants further investigation, as it may be related to edge flow, shear flow, and cell sorting [29, 37–39]. Additionally, studying cell behaviors in monolayers featuring −1 topological defects is also an important direction for future research. Two nonlinear forces can produce outward flow for −1 defects, potentially leading to depletion at the center of these defects.

Our study fills a knowledge gap about how integer topological defects influence collective dynamics in 2D nematic cell sheets system. Defects of +1 topological charge provide aggregation sites, potentially influencing cell transport and morphogenesis in vivo [40]. Furthermore, the identification of emerging nonlinear active forces clarifies our understanding of collective motion in active nematic systems, such as bacterial colonies, cytoskeletal filaments, colloids and 3D morphogenesis [20, 22, 41–44]. We believe our study introduces a new way to quantify and classify cell sheets, highlights the influence of integer topological defects in the biomedical field, and could provide insights for controlling structure formation.

## 4 Methods

### 4.1 Cell culture

Mouse neural progenitor cells were originally from Ryoichiro Kageyama’s lab, and mutants labelled by H2B-mCherry were made by Kyogo Kawaguchi. We used DMEM/F-12, HEPES, no phenol red (Thermo Fisher, 11039021) as medium base supplemented with basic fibroblast growth factor (bFGF) (20 ng/mL, Wako, 060-04543), epidermal growth factor (EGF) (20 ng/mL, Thermo Fisher, 53003-018) and N-2 Max (100x, *R&D* Systems, AR009). 35mm dishes were pre-coated with 10x diluted Matrigel (Corning, 356231) and medium (finally 30*μ*L Matrigel in 1mL medium) overnight in 37^°^*C*. Then we removed the coating agent and placed NPCs uniformly with 2mL medium. The initial cell density is around 5.5 * 10^*−*4^*/μm*^2^. The cultures were incubated at 37^°^*C* with 5% *CO*_2_ at saturated humidity. StemPro Accutase (Gibco, A1110501) was used to disassociate cells during passage progress, and Stem-Cellbanker (ZENOAQ, 11922) was applied for making frozen stocks.

### 4.2 PDMS substrate preparation

We designed the patterns in AutoCAD and produced the chromium mask accordingly, followed by lithography procedures. By spin-coating of SU-8 2002 (MicroChem) at 3000 rpm, we generated 1.2*μm* slots on the silicon wafer after lithography. PDMS liquid with a curing agent proportion of 1/8 (Dow Corning, Sylgard 184) was poured onto the silicon wafer and baked at 70^°^*C* for 3 hours. After solidification, we peeled the PDMS sheet off the silicon wafer and finally obtained a PDMS substrate with ridges. Given that the typical width of an elongated NPC is approximately 10 *μm*, in order to gently guide cells and not hinder their movement too much, we fabricated ridges of width 9*μm* and height 1.2*μm*, and set the distance between ridges to be in the range of 30 − 120*μm*. Multiple types of defects were integrated into a single PDMS sheet, facilitating simultaneous comparisons. Before seeding cells on the PDMS sheet, we applied plasma cleaner to improve hydrophilicity of the surface, sterilized it with 75% ethanol and UV for 30 minutes separately, and washed using PBS and DMEM/F-12. In addition to the micropatterns presented in the main text, we also tested cell behavior on targets with *R*_*max*_ = 300, 600, 900*μm*. The consistent accumulation and the tendency of inward flow remained unchanged.

### 4.3 Imaging

For long time-lapse experiments, NPCs were plated onto Matrigel-coated PDMS substrates in glass base dishes (Iwaki). NPCs were initially cultured in a *CO*_2_ incubator for two days until reaching the desired confluent density, after which they were observed through phase contrast and fluorescent channels of a Leica DMi8 microscope. Tokaihit incubator, which can maintain temperature and humidity and *CO*_2_ concentration, was assembled on the microscope to ensure normal cell growth during recording. Double-channel images were captured every 6 minutes to reduce phototoxicity. Steady integer topological defects induced by the guiding patterns could be observed over several days.

### 4.4 Nematic order

To characterize the local alignment of cells, we measured the orientational field from phase contrast images. Firstly, we computed the gradient of the bright field *I*_*ij*_, where *ij* indicated the positions of image pixels. This brightness gradient was then smoothed using a Gaussian filter with a standard deviation of *σ* = 21*μm*. Subsequently, for each pixel, we built the structure tensor [16, 45],

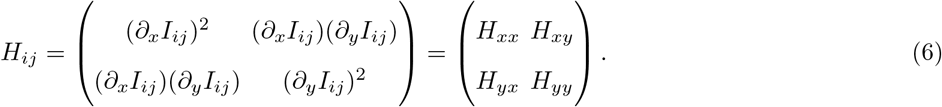

The local orientation *θ*_*ij*_ and coherency *C*_*ij*_ at each pixel was then determined using the formula:

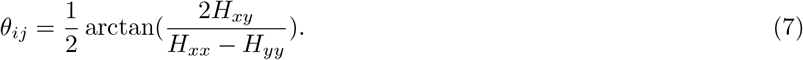

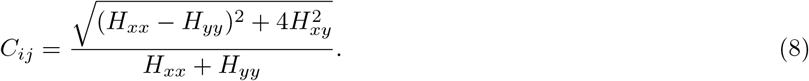

In general, a coherency close to 1 means that the structure is locally 1D, a coherency close to 0 means that there is no preferred direction. The nematic tensor order parameter *Q*_*ij*_ was calculated by

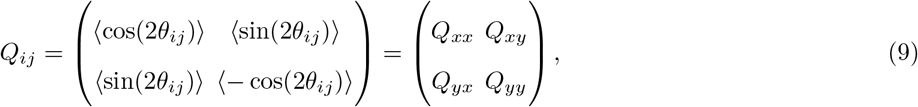

where ⟨·⟩ represents a spatial average calculated using a 2D Gaussian filter with standard deviation *σ* = 42*μm*. Scalar order parameter *S* is defined by 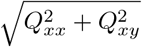. The local spatial average orientational field *θ* was obtained from 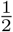 arctan(*Q*_*xy*_*/Q*_*xx*_). For the ideal orientations, *θ*_*ij*_ = *ϕ*_*ij*_ + *θ*_0_ in equation (9), ideal *S* is then obtained by the above steps.

We also extracted orientation fields from cell nuclei in the fluorescent images. Briefly, we employed marker-controlled watershed segmentation to detect the shapes of nuclei. Then we used ellipses to fit shapes and defined the long axis as the orientation of single cell. See Supplementary Information (Section 2) for further detail. At last, these orientations were averaged within a 42 *μm* wide box to derive the local spatial average orientational fields.

### 4.5 Cell velocity measurements

To obtain the cellular velocity, cell nuclei were tracked by TrackMate [46], a ImageJ plugin [47], in the fluorescent images. The LoG detector and simple LAP tracking algorithm were employed. Prior to tracking, the images were stabilized using the Image Stabilizer plugin in ImageJ. For *i*-th trajectory ***r***_*i*_(*t*), instantaneous velocity ***v***_*i*_(*t*) was estimated as the slope determined by {***r***_*i*_(*t* − 2), ***r***_*i*_(*t* − 1), ***r***_*i*_(*t*), ***r***_*i*_(*t* + 1), ***r***_*i*_(*t* + 2)}. The net velocity field ***v***(***x, y***) was obtained by temporally and spatially vector averaging within a 10 *μm* wide box, after which a 2D Gaussian filter with standard deviation *σ* = 42*μm* was applied.

To double-check the angle distributions of velocity fields, we also measured the cellular flows by applying an optical flow algorithm on the fluorescent images. This algorithm is rooted in the Lucas–Kanade derivative of the Gaussian method [48]. Spatial average was done with a 42 *μm* wide box.

In the calculation of autocorrelation of the direction of instantaneous velocity, 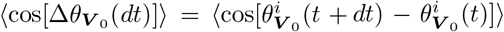, where ***V*** _0_ denotes instantaneous velocity, 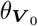 is the direction of ***V*** _0_, *i* is the trajectory number, *t* ∈ [0, duration of *i* − th trajectory], *dt* is time interval (*dt* ∈ [0, duration − *t*]), and ⟨·⟩ is the ensemble average over all the trajectories (*i*) and time (*t*). 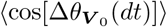 over *dt* can be fitted with exponential exp(−2*dt/τ*_*f*_), where *τ*_*f*_ is the reversal time scale. In the Fig. 3c, *k* = −2*/τ*_*f*_.

### 4.6 Cell density

For cell density quantification, we used the sum of fluorescent images after subtracting background. The background of fluorescent signal was calculated as the average value of the lowest 0.02% of the total pixels. See Supplementary Information (Section 3) for further detail about the transformations between fluorescent intensity and cell number.

### 4.7 Estimation of parameters

For fitting, we normalized *ρ* by the mean density *ρ*_0_. Then the equation (5) was transfromed into

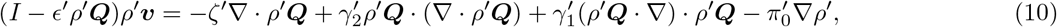

where 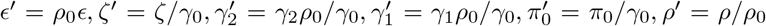. For the convenience in writtng, we set 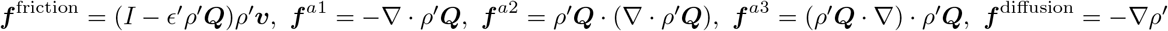. Then we defined parameter vector ***P*** and observation vector ***V***

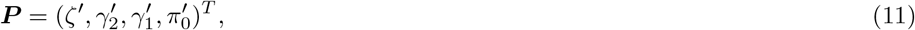

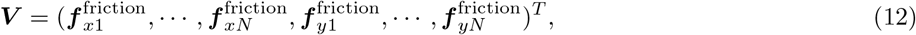

where the subscript *i* = 1, 2, · · ·, *N* denotes the observation at each grid point *i* of fields, *N* is the total number of the observation points, *x, y* denote components in cartesian coordinates. The input matrix ***X*** is defined by

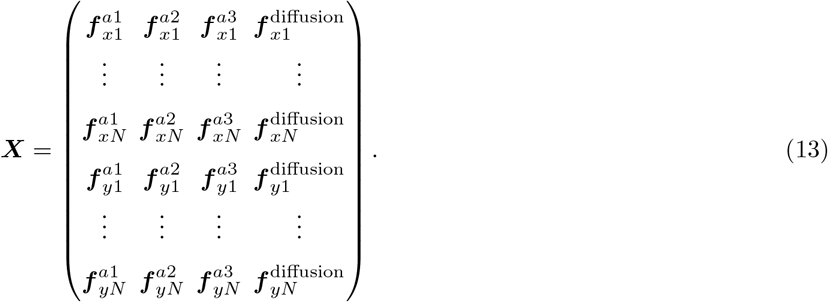

Then equation (9) can be written as ***V*** = ***XP***.

Given that both sides of the equation contain unknown parameters, we considered a fixed *ϵ*^*′*^ at first. We employed Bayesian inference method to keep the diffusion parameter 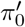 positive [32, 49, 50]. This approach reformulated the equation into the problem of maximizing the posterior probability distribution function Π(***P*** |***V***) of the parameter set for the given ***V*** :

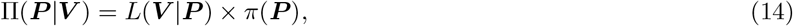

where the likelihood function *L*(***V*** |***P***) was defined as

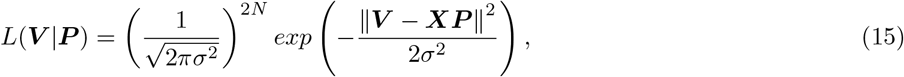

and the prior distribution of the parameter set *π*(***P***) was defined as

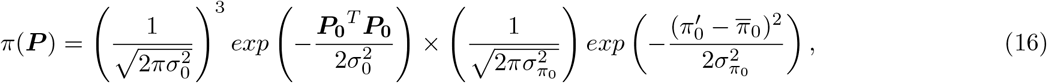

where the parameter vector was separated into two parts, 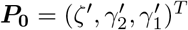 and 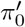. Here, we set a positive bias 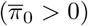 to 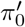 and assumed a Gaussian prior distribution. By maximizing the likelihood function, the optimal parameter set was obtained as

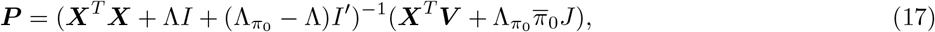

where 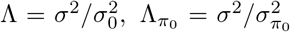, *I* is the identity matrix, *I*^*′*^ is a matrix in which only element *I*^*′*^(4, 4) = 1 and all other elements are 0, *J* = (0, 0, 0, 1)^*T*^. 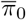 was set by the estimation of the diffusion constant: 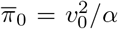 where the instantaneous velocity *v*_0_ = 0.0067*μm/s* and velocity reversal rate *α* = 1*/*3600*s*^*−*1^. The initial values of 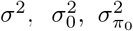 were set as *σ*^2^ = 10^*−*6^(*μm*^2^*/s*^2^), 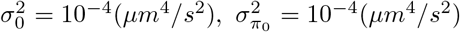. Using the calculated ***P***, *σ*^2^ and 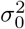 were updated by:

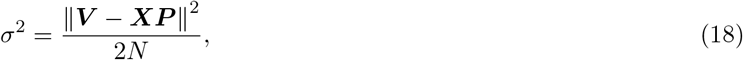

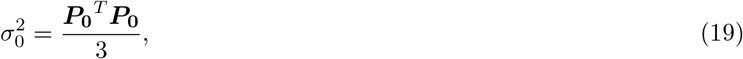

which were subsequently used to update ***P***. This cycle was iterated until the relative change of the Λ per iteration became smaller than 10^*−*3^.

So far, with a fixed *ϵ*^*′*^, we obtained the estimated parameters. Next, we varied *ϵ*^*′*^, and for each *ϵ*^*′*^, we used the Bayesian method to obtain the estimated velocities as stated earlier. Then we calculated the determination coefficients between the estimated velocities and the measured velocities. We selected the optimal *ϵ*^*′*^ and other parameters which maximized the determination coefficients until the relative change of determination coefficients became smaller than 10^*−*5^.

### 4.8 Correction for the predicted outward flow of the center

Slight outward flows appear at centers of defects with large |*θ*_0_|, where the density is much higher (Fig. 4g, Extended Data Fig. 8a,b). Here, we neglect the ∇*ρ* term in equation (5) around centers (*r < R*_*min*_) according to an assumption that cells can readily move to the third dimension and form cell mounds at high densities under compression. Additionally, according to equation (3,4), active forces near centers are sensitive to the amplitude and gradient of *S* which we could not reliably measure due to equipment limitations. By using ideal orientations (*θ* = *θ*_*p*_) and ideal *S* (Extended Data Fig. 9b) according to the definition of defects and setting *π*_0_^*′*^ = 0 at regions with *r <* 80*μm* (with other parameters as best-fit values), the outward cellular flows vanish basically (Extended Data Fig. 9a,c). This method causes fluctuations in net velocity amplitudes around *r* = 80*μm* (Extended Data Fig. 9a,c). To address this, we employ linear interpolation to replace the fluctuating values (Extended Data Fig. 9e).

## Code and data availability

All the codes and the data that support the findings of this study are available from the Z.Z. or M.S. upon request.

## Supplementary information

Supplementary information is available in the online version of the paper.

## Acknowledgements

We are grateful to Jacques Prost, Sriram Ramaswamy, Xiaqing Shi, Carles Blanch-Mercader, Aurelio Patelli, Eric Bertin, and Benoît Mahault for for insightful discussions. We thank Kirsten D. Endresen for making micropatterns at the initial stage. M.S. acknowledges support from the Research Fund for International Scientists of NFSC (12174254,12250710131). H.Z. acknowledges support from NSFC (12225410 and 12074243).

## Author contributions

M.S., F.S., H.Z., and Z.Z. conceived and designed the research. Z.Z., H.L., and Y.Y. performed the experiments. Z.Z. analyzed the experimental data. K.K. provided materials and tools. M.S., Y.Z., and H.C. developed the theoretical part. All the authors participated in writing and revising the manuscript.

---

**Extended Data Fig. 1.**
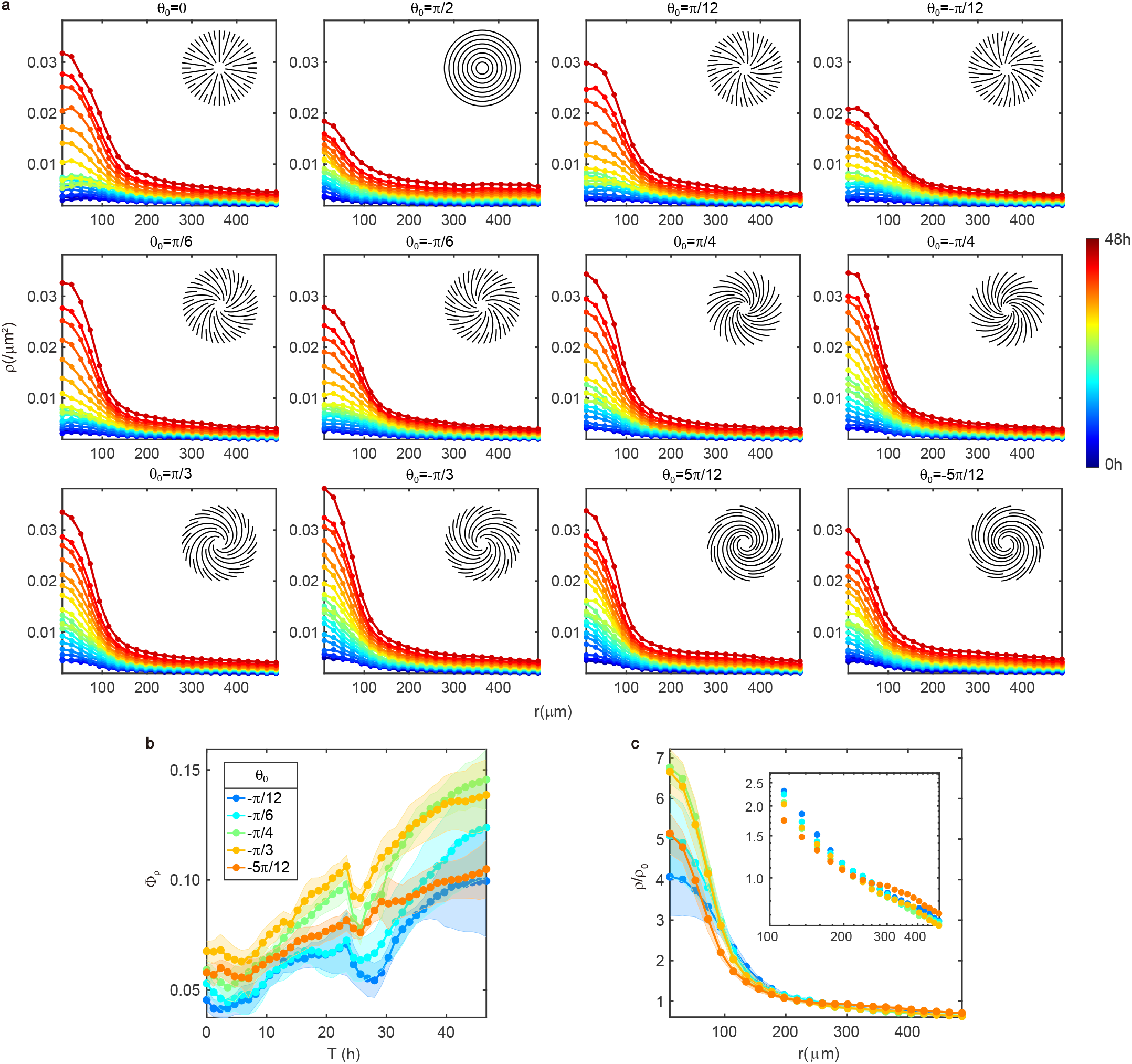
Cell density evolution of all defects which are arranged on one PDMS sheet. **a**, Time evolution of cell density distributions. Density profiles are obtained by taking the average of data captured every 1 hour, and the color encodes time lapse. **b**, Average percentage of cells Φ_*ρ*_ (Φ_*ρ*_ = *n*(*r < R*_*min*_)*/n*(*r < R*_*max*_), where n is the cell number) assemblies with time on defects with negative *θ*_0_. Colors of curves depict different defects. **c**, Normalized radial density profiles by mean density *ρ*_0_ when *ρ*_0_ = 0.009*/μm*^2^. Inset: A log-log plot for the region with *r >* 100*μm*. Legend in **b**. N of each type of defects is 6.

**Extended Data Fig. 2.**
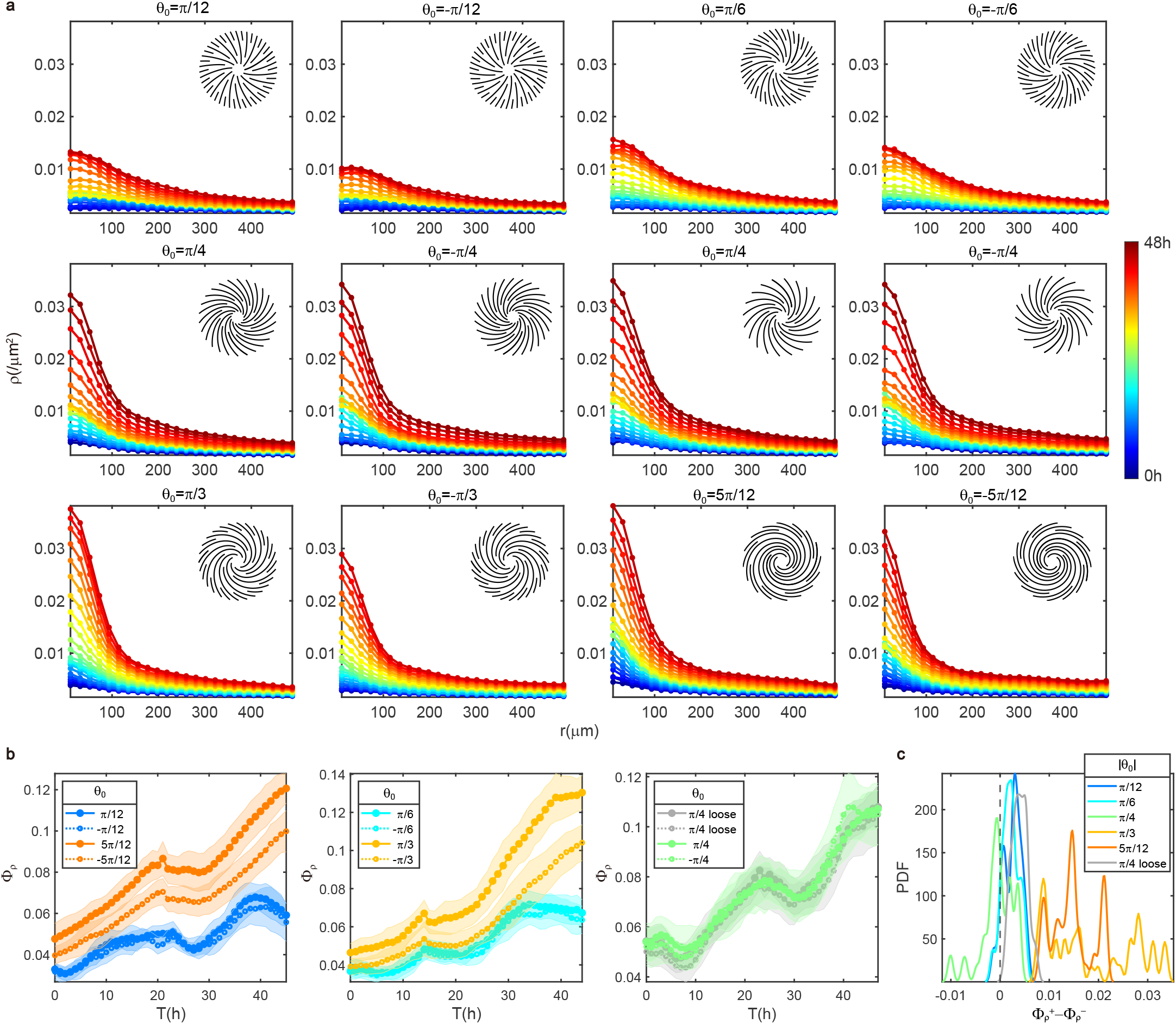
NPCs accumulate at the center of spirals and show more prominent accumulation on counterclockwise spirals compared with clockwise spirals. **a**, Time evolution of cell density distributions on spirals. Density profiles are obtained by taking average of data captured every 1 hour, and the color encodes time lapse. Defects with *θ*_0_ = ±*π/*12, ±5*π/*12 are on one PDMS sheet, so are the defects with *θ*_0_ = ±*π/*6, ±*π/*3, and the defects with *θ*_0_ = ±*π/*4, ±*π/*4 (looser confinements). N of each type of spirals is 16. **b**, Average percentage of cells Φ_*ρ*_ (Φ_*ρ*_ = *n*(*r < R*_*min*_)*/n*(*r < R*_*max*_), where n is the cell number) assemblies with time on defects with negative *θ*_0_ and on patterns with looser confinements. Colors of curves depict different defects. **c**, Distribution of the difference between Φ_*ρ*_ of clockwise spirals and counterclockwise spirals. The difference is calculated every 1 hour for defects with opposite sign but the same absolute value of *θ*_0_. In **b** and **c**, data are presented as mean ± s.d.

**Extended Data Fig. 3.**
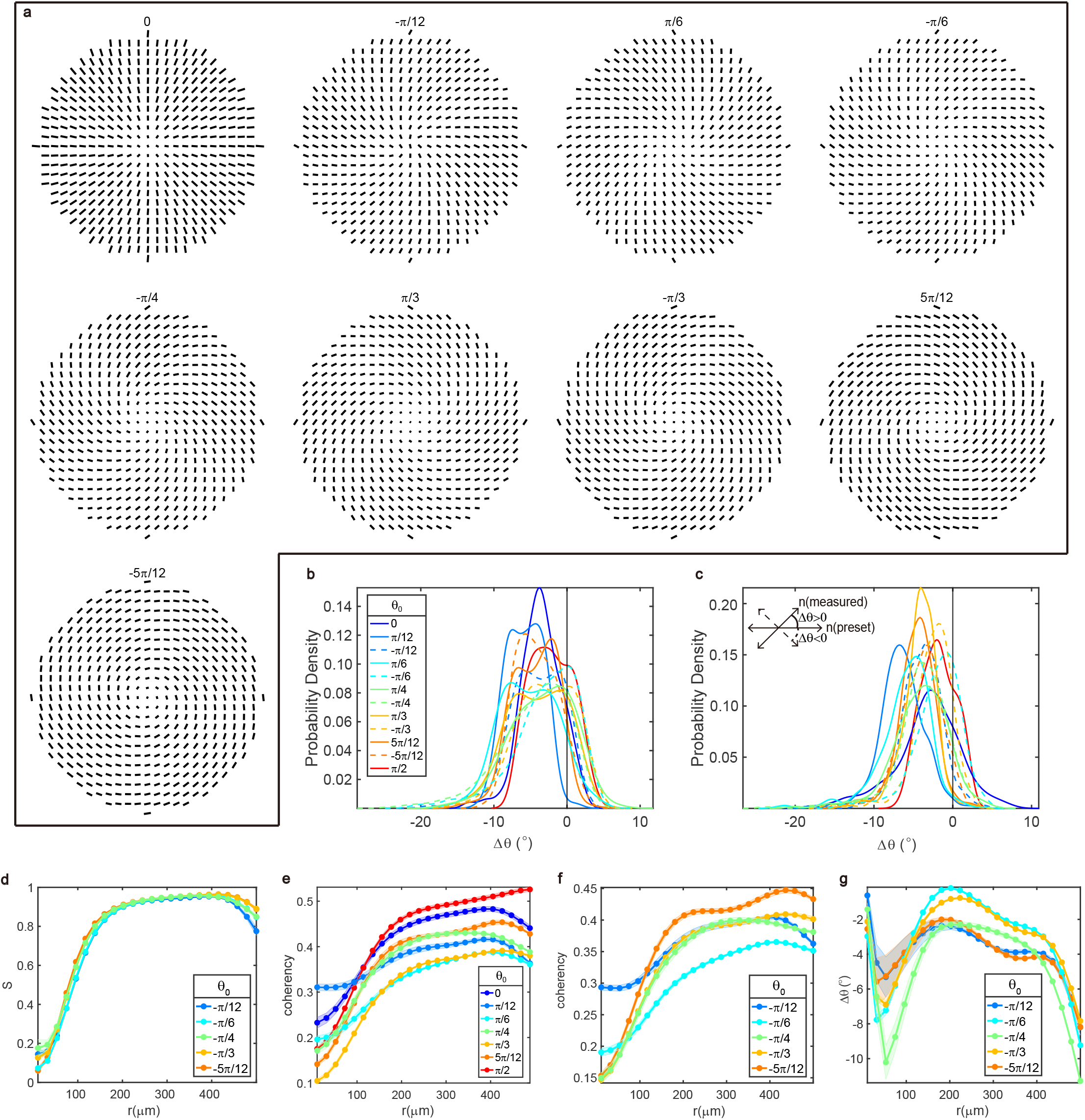
Orientational analysis on aster, spirals, and target. **a**, Measured orientational fields by structure tensor method. The length of line corresponds to the coherency. Titles are values of *θ*_0_. **b**, Probability density function of the angle (Δ*θ*) between the experimentally measured cell orientation by structure tensor method and orientation preset by PDMS pattern. **c**, Probability density function of the angle (Δ*θ*) between the experimentally measured cell orientation by detecting single cell and orientation preset by PDMS pattern. Inset shows the definition of Δ*θ*. Legend in **b**. In **b, c**, *r <* 100*μm* was excluded. **d**, Radial profiles of order parameter S for clockwise spirals. **e**, Radial profiles of coherency for aster, counterclockwise spirals and target. **f**, Same as **d** but for coherency. **g**, Same as **d** but for Δ*θ* detected by structure tensor method. In **d, g**, data are presented as mean ± s.d., and s.d. are calculated from different times. In **e, f**, data are presented as mean ± s.e.m., and s.e.m. are calculated from different times. In **a**−**g**, data are averaged by 32 asters (10h), 16 spirals for each type (10h) and 32 targets (10h).

**Extended Data Fig. 4.**
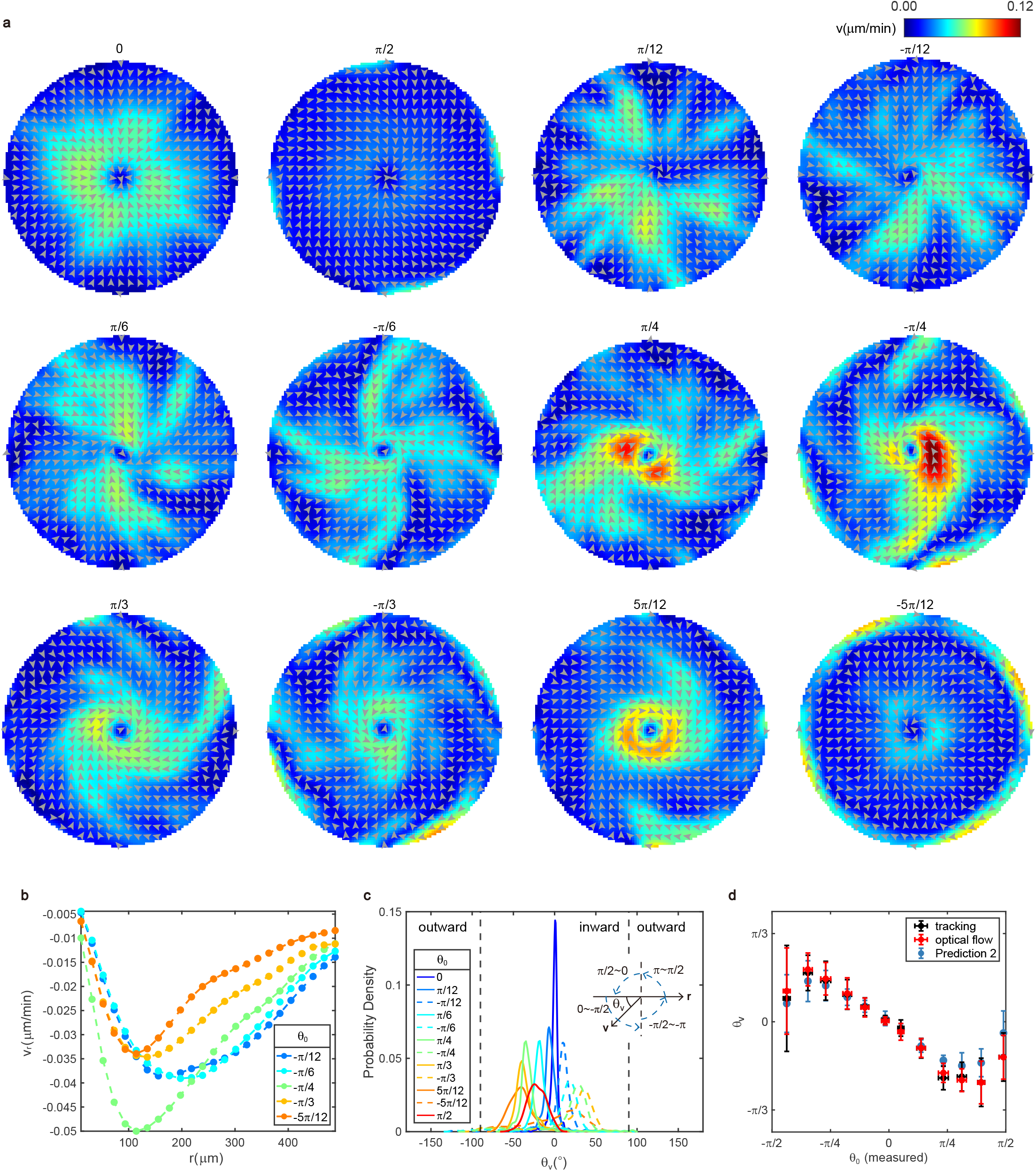
Net velocity fields on aster, spirals, and target. **a**, Net velocity fields calculated from tracking results across all the defects. Colors depict the amplitude of net velocity. Titles are values of *θ*_0_. **b**, Radial profiles of the net radial velocity for clockwise spirals. **c**, Probability density function of the angle (*θ*_*v*_) between net velocity and inward radial direction on all types of defects. Velocity fields are calculated from optical flow method. Insets show the definition of *θ*_*v*_. Regions with *r <* 100*μm* are excluded. **d**, Measured and predicted *θ*_*v*_ versus measured *θ*_0_. Black dots are the same experimental data as those in Fig. 2**i**. *θ*_*v*_ of red dots are obtained from the location of peaks in **c**. Prediction 2 comes from method (ii). In **a** − **d**, data are averaged by 32 asters (10h), 16 spirals for each type (10h) and 32 targets (10h).

**Extended Data Fig. 5.**
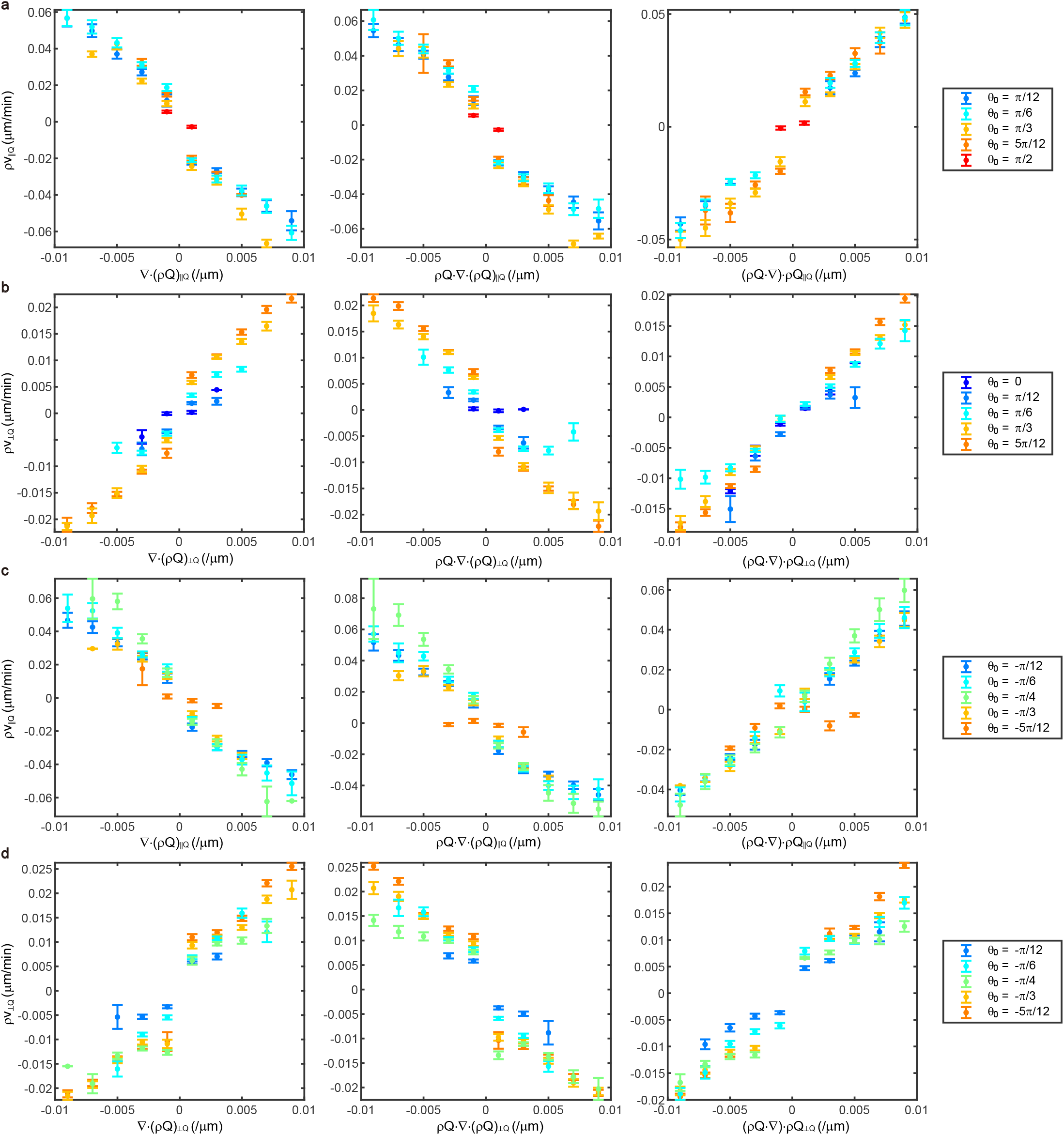
The comparison between components of cellular flow *ρv* and components of active forces. ∥ ***Q*** (⊥ ***Q***) is defined as the component parallel (perpendicular) to the principal axis of ***Q*. a**, Comparisons between parallel components for defects with positive *θ*_0_. **b**, Comparisons between perpendicular components for the aster and spirals with positive *θ*_0_. **c**, Comparisons between parallel components for spirals with negative *θ*_0_. **d**, Comparisons between perpendicular components for spirals with negative *θ*_0_. Data are presented as mean ± s.e.m.

**Extended Data Fig. 6.**
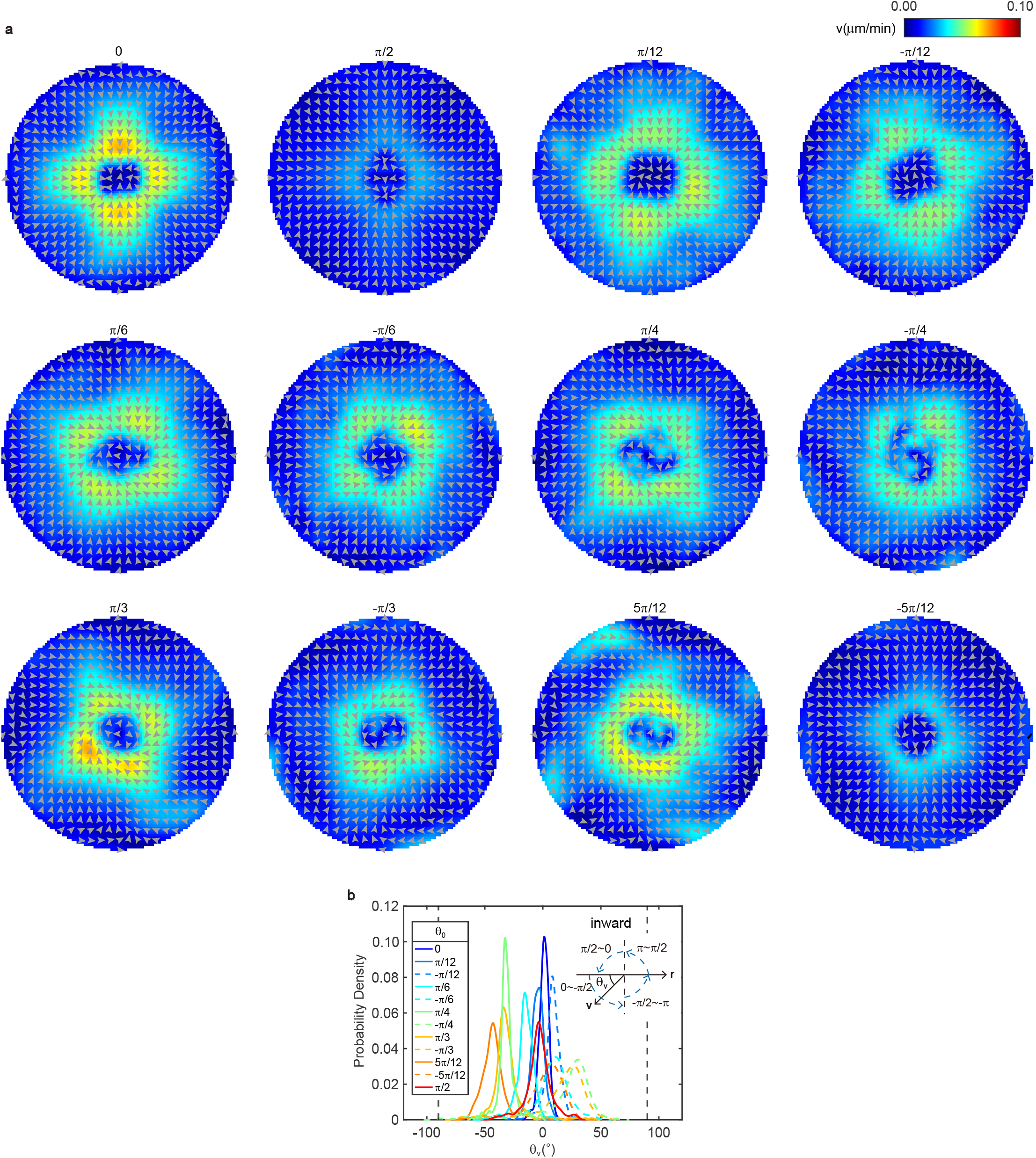
Predicted velocities by best-fit parameters on aster, spirals, and target. **a**, Predicted net velocity fields across all the defects. Colors depict the amplitude of net velocity. Titles are values of *θ*_0_. **b**, Probability density function of the angle (*θ*_*v*_) between predicted net velocities and inward radial direction on all types of defects. Insets show the definition of *θ*_*v*_. The prediction method is method (i). Regions with *r <* 100*μm* are excluded.

**Extended Data Fig. 7.**
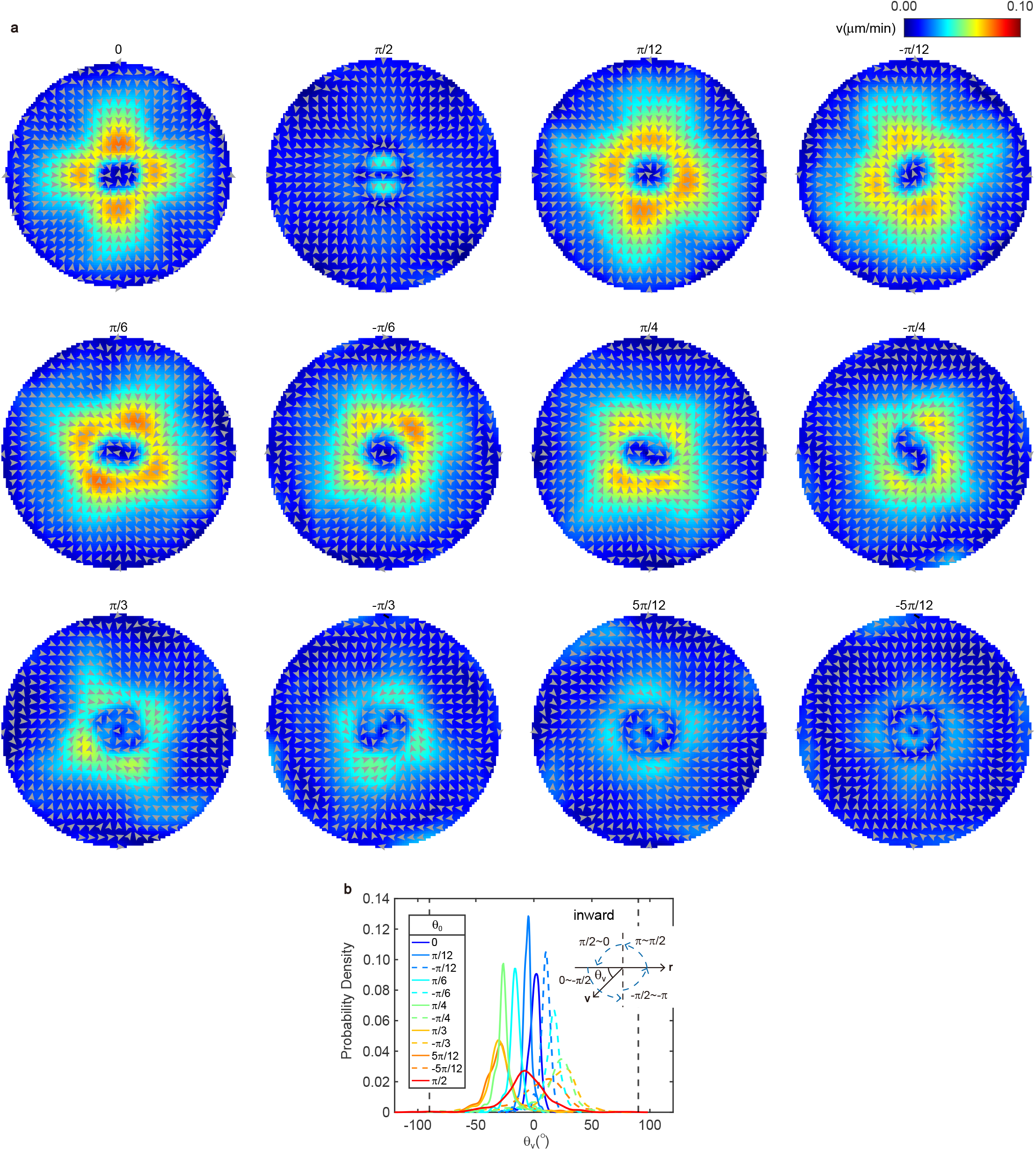
Predicted velocities by average parameters on aster, spirals, and target. **a**, Predicted net velocity fields across all the defects. Colors depict the amplitude of net velocity. Titles are values of *θ*_0_. **b**, Probability density function of the angle (*θ*_*v*_) between predicted net velocities and inward radial direction on all types of defects. Insets show the definition of *θ*_*v*_. The prediction method is method (ii). Regions with *r <* 100*μm* are excluded.

**Extended Data Fig. 8.**
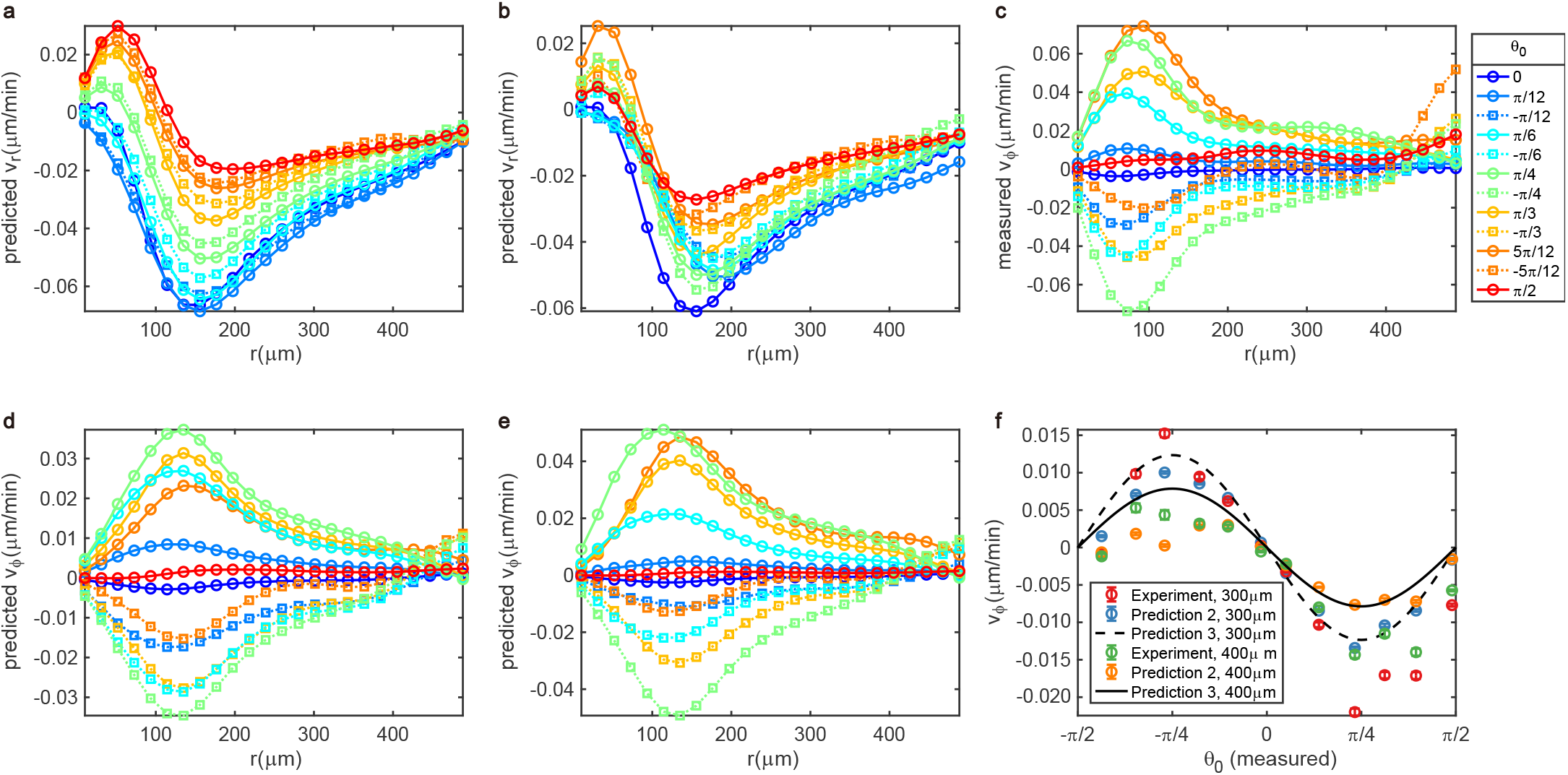
Predicted and measured velocity components on aster, spirals, and target. **a, b**, Radial profiles of the predicted radial velocities by method (ii) (**a**) and method (i) (**b**). Legend in **c. c**, Radial profiles of the measured azimuthal velocities (*v*_*ϕ*_). **d, e**, Radial profiles of the predicted azimuthal velocities by method (ii) (**d**) and method (i) (**e**). Legend in **c. f**, Comparison between measured azimuthal velocities and predictions at *r* = 300*μm* and *r* = 400*μm* of all the defects. Method of prediction 2 (3) is method (ii) ((iii)). Data are presented as mean ± s.e.m.

**Extended Data Fig. 9.**
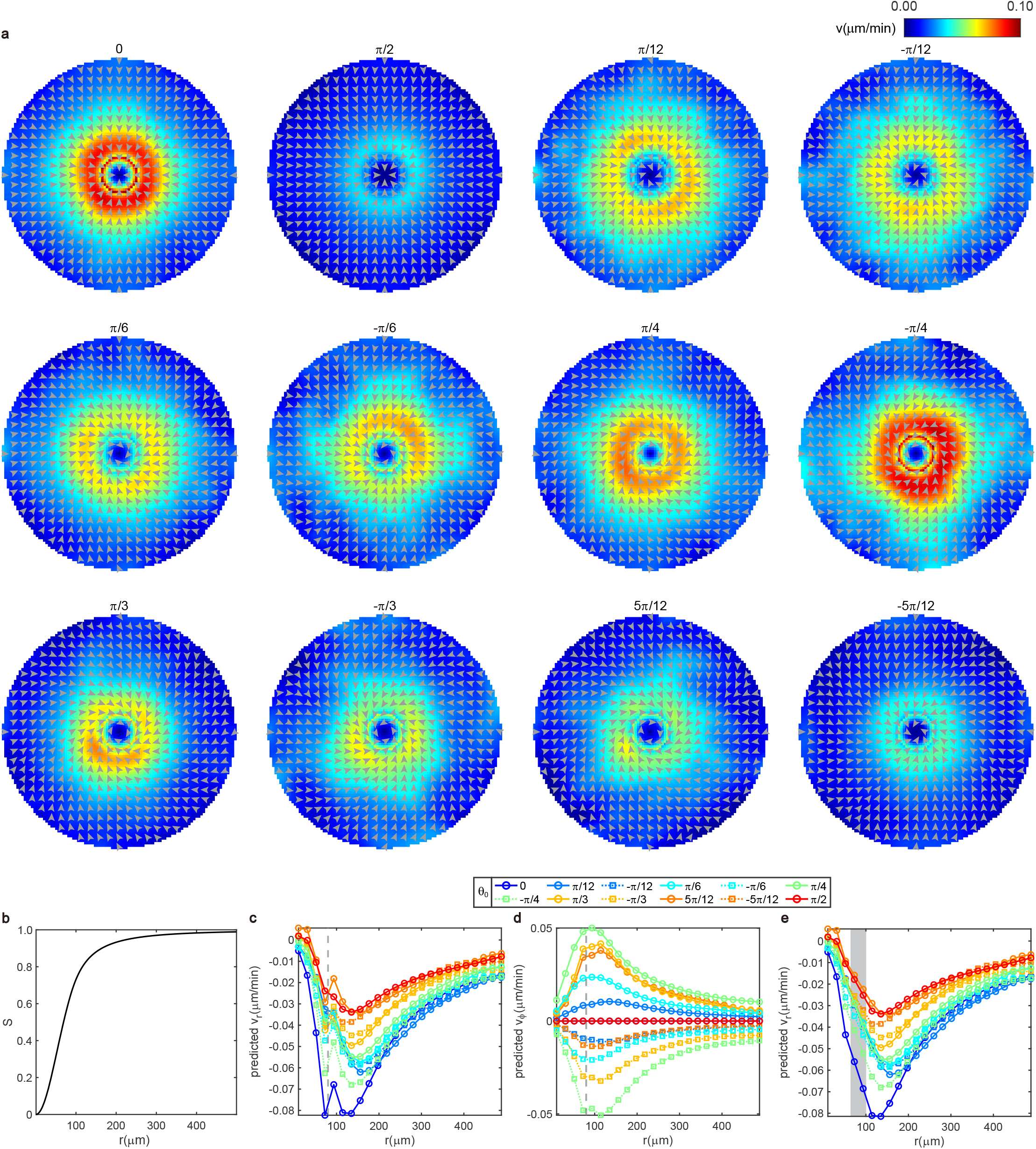
Correction for the predicted outward flow of the center. **a**, Predicted velocities by best-fit parameters and perfect orientations according to the definition of integer topological defects on aster, spirals, and target. In regions with *r <* 80*μm*, 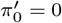. Titles are values of *θ*_0_. **b**, Radial profiles of order parameter *S* on the ideal integer topological defects. **c, d, e**, Radial profiles of the predicted radial velocities (**c, e**), and azimuthal velocities (**d**). In regions with *r <* 80*μm* (Left part of the dashed line in **c, d**), 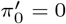. In **e**, we use linear interpolation to reduce fluctuations around *r* = 80*μm* (gray region).

**Extended Data Table 1.** Best-fit parameters and determinate coefficients (*R*^2^) for each kind of pattern.

| $\theta_0$ | Parameters | | | | | $R^2$ |
| --- | --- | --- | --- | --- | --- | --- |
| | $\zeta'(\mu m^2/s)$ | $\gamma_2'(\mu m^2/s)$ | $\gamma_1'(\mu m^2/s)$ | $\pi_0'(\mu m^2/s)$ | $\epsilon'$ | |
| $-5\pi/12$ | 0.0197 | -0.0457 | 0.0316 | 0.0403 | 0.1300 | 0.8224 |
| $-\pi/3$ | 0.0264 | -0.0379 | 0.0498 | 0.0689 | 0.1900 | 0.8371 |
| $-\pi/4$ | 0.0375 | -0.0327 | 0.0648 | 0.1041 | 0.2200 | 0.8519 |
| $-\pi/6$ | 0.0268 | -0.0208 | 0.0724 | 0.1166 | 0.0210 | 0.8284 |
| $-\pi/12$ | 0.0164 | -0.0079 | 0.0656 | 0.0991 | 0.2000 | 0.8756 |
| 0 | 0.0275 | -0.0152 | 0.0535 | 0.0594 | 0.1500 | 0.9551 |
| $\pi/12$ | 0.0151 | -0.0043 | 0.0758 | 0.0994 | 0.1800 | 0.8876 |
| $\pi/6$ | 0.0296 | -0.0069 | 0.0679 | 0.0820 | 0.0580 | 0.8740 |
| $\pi/4$ | 0.0590 | -0.0230 | 0.0557 | 0.0512 | 0.0200 | 0.8915 |
| $\pi/3$ | 0.0321 | -0.0296 | 0.0507 | 0.0503 | 0.2010 | 0.9060 |
| $5\pi/12$ | 0.0434 | -0.0508 | 0.0579 | 0.0862 | 0.3110 | 0.8928 |
| $\pi/2$ | 0.0289 | -0.0257 | 0.0564 | 0.0732 | 0.1476 | 0.8173 |

