## Supplementary Information for "Integer topological defects provide a new way to quantify and classify cell sheets"

#### 1 Supplementary videos

**Movie S1**: NPCs move along or cross over the ridges.

**Movie S2**: Time-lapse fluorescent images of all +1 defects on one PDMS sheet.

**Movie S3**: Time evolution of instantaneous velocity, net velocity and density field on the aster during the steady accumulation periods.

**Movie S4**: Time evolution of instantaneous velocity, net velocity and density field on the target during the steady accumulation periods.

**Movie S5**: Time evolution of instantaneous velocity, net velocity and density field on the counterclockwise spiral ($\theta_0 = \pi/4$) during the steady accumulation periods.

**Movie S6**: Time evolution of instantaneous velocity, net velocity and density field on the clockwise spiral ($\theta_0 = -\pi/4$) during the steady accumulation periods.

### 2 Measurement of orientational field from cell nuclei

As described in Methods, we employed marker-controlled watershed segmentation to detect the shapes of nuclei [1]. We first applied a top-hat transformation and bilateral filtering to images. Then we utilized extended maxima transform to identify internal markers inside cell nuclei. Next, we computed the watershed transform of the distance transform of the internal markers to identify the external markers in the background. These internal and external markers were then utilized to modify the gradient image using a procedure known as minima imposition so that regional minima occurred only in marked locations. By computing the watershed transform of the marker-modified gradient images, we obtained the contours of cell nuclei. After filling the contours and removing small objects, we used regionprops function in MATLAB to determine the orientations of nuclei. In Fig. S1, we show shapes and orientations of some cells detected by this method.

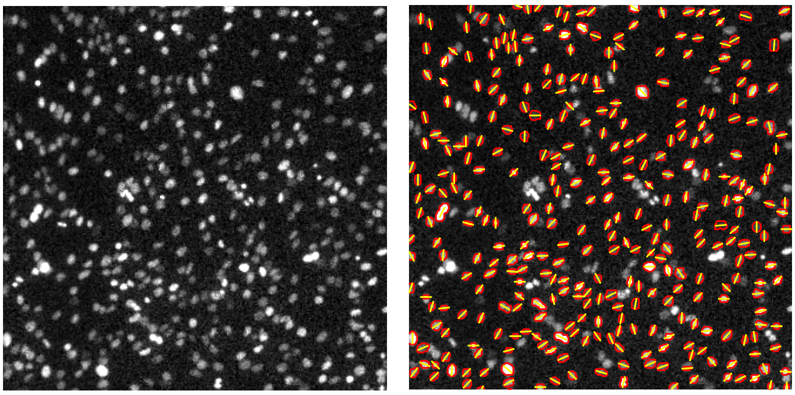

**Fig. S1** Schematic diagram of detecting cell orientations from fluorescent images. Left: original data; Right: cell shapes (red) and orientations (yellow).

### 3 Transformation between fluorescent intensity and cell number

For each frame in the time-lapse fluorescent images, we counted the number of cells using the LoG detector in TrackMate. At low densities, we found a linear relation between fluorescent intensity and cell number from the counts (Fig. S2). Using this linear relation, we can predict cell density from fluorescent intensity when counting becomes imprecise in high-density regions.

### 4 Theory

To understand the mechanism behind the inward cellular flows toward the cores of integer defects, we conducted simulations of a model of interacting active particles under external fields inducing integer defects, derived from it an hydrodynamic theory by the Boltzmann-Ginzburg-Landau approach, and analyzed its solutions. All details can be

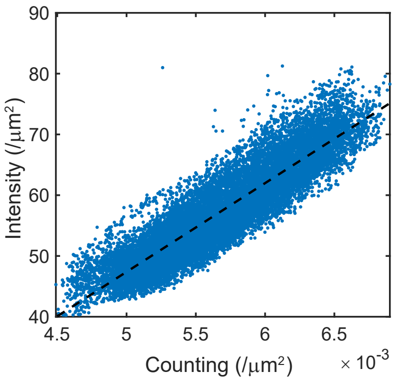


**Fig. S2** The relation between fluorescent intensity and cell density obtained from counting cell nuclei.

found in [2]. The particle-level model and the hydrodynamic theory derived from it both display accumulation toward the core of +1 defects, irrespective of their particular phase $\theta_0$.

#### 4.1 Particle model

Here, we consider the model studied in [3] supplemented by an external field patterned with various +1 defect shapes: point particles move at constant speed $v_0$ along the unit orientation/velocity vector $\boldsymbol{e}_i^t$ which reverses stochastically at rate $k$. Particles align nematically with neighbors within unit distance. They also experience repulsive torques from the same neighbors. The time evolution of positions $\boldsymbol{r}_i^t$ and orientations $\boldsymbol{e}_i^t$ obeys

$$\boldsymbol{r}_i^{t+1} = \boldsymbol{r}_i^t + v_0\, \boldsymbol{e}_i^{t+1}, \tag{1}$$

$$\theta_i^{t+1} = \arg\{\epsilon_t[\langle \mathrm{sgn}(\boldsymbol{e}_i^t \cdot \boldsymbol{e}_j^t)\boldsymbol{e}_j^t\rangle_j + C_p\, \mathrm{sgn}(\boldsymbol{e}_i^t \cdot \boldsymbol{e}_i^p)\boldsymbol{e}_i^p] + C_r\, \langle \hat{r}_{ji}^t\rangle_j\} + \eta\, \chi_i^t, \tag{2}$$

where $\theta_i^t = \arg(\boldsymbol{e}_i^t)$, $\langle\cdot\rangle_j$ denotes the average over neighbors $j$ within a unit distance of $i$ including $i$ for the alignment. $\epsilon_t = \pm 1$ reverses sign with probability $k \in [0, 0.5]$, $C_r$ is the strength of repulsion and $\hat{r}_{ji}^t$ is the unit vector from the particle $j$ to $i$. $\eta$ is the strength of angular noise. $\chi_i^t \in [-\frac{\pi}{2}, \frac{\pi}{2}]$ is a white noise taken from uniform distribution. The external guiding field of strength $C_p$ is expressed by the local unit vector $\boldsymbol{e}_i^p$ which is defined as $\arg(\boldsymbol{e}_i^p) = \theta^p(\boldsymbol{r})\delta(\boldsymbol{r}-\boldsymbol{r}_i^t)$ with a prescribed local angle $\theta^p(\boldsymbol{r})$ to induce the nematic alignment to the particles at each position $\boldsymbol{r}_i^t$, where $\theta^p = \phi + \theta_0$. We performed particle simulations and obtained systematic accumulation on various types of defects.

#### 4.2 Continuous theory

Employing the Boltzmann-Ginzburg-Landau approach to the particle model, we derived the hydrodynamic equations governing the time evolution of density $\rho$, velocity field $\boldsymbol{v}$, and nematic tensor $\boldsymbol{Q}$,

$$\partial_t \rho = -\frac{1}{2} v_0 (\nabla^* f_1 + \nabla f_1^*), \tag{3}$$

$$\partial_t f_1 = (\alpha[\rho] - \beta|f_2|^2) f_1 + \sigma f_1^* f_2 - \pi_0[\rho]\nabla\rho - \zeta[\rho]\nabla^* f_2 - \lambda_n f_2 \nabla^* \rho + \lambda_1 f_1 \nabla^* f_1$$

$$+\lambda_2 f_1 \nabla f_1^* + \lambda_3 f_1^* \nabla f_1 + \gamma_2 f_2 \nabla f_2^* + \gamma_1 f_2^* \nabla f_2 + \frac{C_p}{2} e^{2i\theta_p} f_1^*, \tag{4}$$

$$\partial_t f_2 = F(f_1, f_2) + C_p e^{2i\theta_p} \rho, \tag{5}$$

where $(\mathcal{R}f_1, \mathcal{I}f_1) = \rho \boldsymbol{v}/v_0$, $(\mathcal{R}f_2, \mathcal{I}f_2) = \rho(\boldsymbol{Q}_{xx}, \boldsymbol{Q}_{xy})$. Detailed descriptions of all terms and coefficients can be found in [2]. Simulations confirmed consistent attraction to the cores in some parameter space. Considering that $\boldsymbol{Q}$ is essentially enslaved to the imposed patterns and the small cell speed observed in the experiments, here we neglect the evolution of both the $\boldsymbol{Q}$ and the $\boldsymbol{v}$ fields. Neglecting further anisotropic diffusion terms, equations (3-5) are simplified as follows:

$$\partial_t \rho = -\nabla \cdot (\rho \boldsymbol{v}), \tag{6}$$

$$\gamma_0 (I - \epsilon \rho Q) \rho \boldsymbol{v} = -\overline{\zeta}[\rho] \nabla \cdot \rho Q + \overline{\gamma}_2 \rho Q \cdot (\nabla \cdot \rho Q) + \overline{\gamma}_1 (\rho Q \cdot \nabla) \cdot \rho Q - \overline{\pi}_0[\rho] \nabla \rho, \tag{7}$$

where $\gamma_0 = -(\alpha[\rho] - \beta |\rho Q|^2)$, $\epsilon = \sigma/\gamma_0$. The coefficients for the equation (7) read:

$$\begin{aligned}
\alpha[\rho] &= \rho \frac{4}{\pi} \left( P_1 - \frac{4}{3} \right) + P_1 - 1 - 2a \\
\beta &= \frac{8}{3\pi} \left( P_1 - \frac{2}{7} \right) \frac{\frac{4}{\pi} \left( P_3 + \frac{4}{5} \right)}{\rho_0 \frac{272}{35\pi} - P_3 + 1 + 2a} \\
\sigma &= \frac{16}{5\pi} \\
\overline{\pi}_0[\rho] &= v_0 \pi_0[\rho] = v_0 \left( \frac{v_0}{2} + \frac{b_1}{4} \left( \pi - 2\sqrt{2} P_1 \right) \rho \right) \\
\overline{\zeta}[\rho] &= v_0 \zeta[\rho] = v_0 \left( \frac{v_0}{2} - \frac{b_1}{6} \sqrt{2} P_1 \rho \right) \\
\overline{\gamma}_1 &= v_0 \gamma_1 = v_0 \left( \frac{4}{3\pi} \left( P_1 - \frac{2}{7} \right) \frac{v_0 - \sqrt{2} b_1 P_3 \rho_0}{\rho_0 \frac{272}{35\pi} - P_3 + 1 + 2a} + \frac{b_1}{6} \sqrt{2} P_1 \right), \\
\overline{\gamma}_2 &= v_0 \gamma_2 = v_0 \left( -\frac{b_1}{10} \sqrt{2} P_1 \right),
\end{aligned}$$

where $b_1$ is related to the strength $C_r$ of the repulsive torques in the particle-level model, $P_k$ are the Fourier modes of the noise distribution, $a$ is the reversal rate. For simplicity, $\zeta, \gamma_2, \gamma_1, \pi_0, \lambda_n$ in the main text refer to $\overline{\zeta}, \overline{\gamma}_2, \overline{\gamma}_1, \overline{\pi}_0, \overline{\lambda}_n$ here. In the experimental data, cell speed and density profiles do not change much outside the defect core region. We also assume that repulsion and noise remain relatively constant. Therefore, when fitting the data from regions outside the defect centers, we treat these parameters as constants.

#### 4.3 Velocity field around integer topological defects

In the main text, we present the forms of linear and nonlinear active forces around integer topological defects. By solving equation (5) of the main text, we obtain the net velocity field ($\boldsymbol{v}^a$) generated by the active force terms. For the ideal cases, $\theta = \theta_p = \phi + \theta_0$, the components of $\boldsymbol{v}^a$ in polar coordinates are

$$\boldsymbol{v}_r^a = -\frac{\zeta}{\gamma_0}\frac{\cos(2\theta_0)+\epsilon\rho S}{1-\epsilon^2\rho^2S^2}\left[\frac{2\rho S}{r}+(\rho S)'\right]+\frac{\rho S}{\gamma_0}\frac{1+\epsilon\rho S\cos(2\theta_0)}{1-\epsilon^2\rho^2S^2}\left\{\gamma_2\left[\frac{2\rho S}{r}+(\rho S)'\right]+\gamma_1\left[(\rho S)'-\frac{2\rho S}{r}\right]\right\}, \quad (8)$$

$$\boldsymbol{v}_\phi^a = -\frac{\zeta}{\gamma_0}\frac{\sin(2\theta_0)}{1-\epsilon^2\rho^2S^2}\left[\frac{2\rho S}{r}+(\rho S)'\right]+\frac{\rho S}{\gamma_0}\frac{\epsilon\rho S\sin(2\theta_0)}{1-\epsilon^2\rho^2S^2}\left\{\gamma_2\left[\frac{2\rho S}{r}+(\rho S)'\right]+\gamma_1\left[(\rho S)'-\frac{2\rho S}{r}\right]\right\}, \quad (9)$$

where $(\rho S)' = \frac{d(\rho(r)S(r))}{dr}$.
